# Human regulatory T cells at the maternal-fetal interface show functional site-specific adaptation with tumor-infiltrating-like features

**DOI:** 10.1101/820753

**Authors:** Judith Wienke, Laura Brouwers, Leone van der Burg, Michal Mokry, Rianne C. Scholman, Peter G.J. Nikkels, Bas van Rijn, Femke van Wijk

## Abstract

**Objectives:** Regulatory T cells (Tregs) are crucial for maintaining immune tolerance against the semi-allogeneic fetus during pregnancy. Since their functional profile at the human maternal-fetal interface is still elusive, we investigated the transcriptional profile and functional adaptation of human uterine Tregs (uTregs) during pregnancy.

**Methods:** Blood and uterine biopsies from the placental bed (=maternal-fetal interface) and incision site (=control), were obtained from women with uneventful pregnancies undergoing primary Caesarean section. Tregs and CD4^+^ non-Tregs (Tconv) were isolated for transcriptomic profiling by Cel-Seq2. Results were validated on protein and single cell level by flow cytometry.

**Results:** Placental bed uterine Tregs (uTregs) showed elevated expression of Treg signature markers compared to blood Tregs, including FOXP3, CTLA4 and TIGIT. The uTreg transcriptional profile was indicative of late-stage effector Treg differentiation and chronic activation with high expression of immune checkpoints GITR, TNFR2, OX-40, 4-1BB, genes associated with suppressive capacity (CTLA4, HAVCR2, IL10, IL2RA, LAYN, PDCD1), activation (HLA-DR, LRRC32), and transcription factors MAF, PRDM1, BATF, and VDR. uTregs mirrored uTconv Th1 polarization, and characteristics indicating tissue-residency, including high CD69, CCR1, and CXCR6. The particular transcriptional signature of placental bed uTregs overlapped strongly with the specialized profile of human tumor-infiltrating Tregs, and, remarkably, was more pronounced at the placental bed than uterine control site.

**Conclusion:** uTregs at the maternal-fetal interface acquire a highly differentiated effector Treg profile similar to tumor-infiltrating Tregs, which is locally enriched compared to a distant uterine site. This introduces the novel concept of site-specific transcriptional adaptation of human Tregs within one organ.

## INTRODUCTION

In the past decade, T cells have been identified in a spectrum of human and murine non-lymphoid tissues.^1,2^ T cells residing in tissues for prolonged periods are thought to serve as first-line responders to infections.^1,2^ These tissue-resident memory T cells (TRM) do not recirculate and are characterized by expression of signature molecules such CD69, which prevents their tissue egress.^1,3–8^ TRM adapt to their tissue environments by acquiring a specialized functional phenotype that depends on micro-environmental cues.^9,10^ Also regulatory T cells (Tregs), which are critical gatekeepers of immune homeostasis,^11^ have been recently identified in various murine and human tissues.^12–17^ Like TRM, Tregs can become resident and adapt to their microenvironment.^12,14,15,18–20^ Tissue-resident Tregs gain a polarized phenotype compared to circulating Tregs, with functional specialization depending on the tissue or organ, which is controlled on a transcriptional level.^12,14,16,21,22^ Although increasing evidence in mice supports this functional adaptation of Tregs to non-lymphoid tissue environments,^23^ studies investigating tissue adaptation of human Treg are still scarce. Only recently, the first transcriptional profiles of Tregs in healthy human tissues were published.^14,15^ A human tissue environment in which transcriptional adaptation of Tregs has gained interest due to the important therapeutic implications, is the tumor environment.^24^ Tumor-infiltrating Tregs (TITR) have been shown to display a unique and specialized transcriptional signature,^25^ which is associated with activation and functional specialization of Tregs at these sites, including increased suppressive capacity.^25–27^ In tissues, and especially in tumors, Tregs undergo differentiation reminiscent of effector Tregs. Effector Tregs are a subset of Tregs with potent suppressive capacity, which are characterized by expression of CD45RO, and increased CD25, CTLA-4, and HLA-DR.^28–31^ Furthermore, effector Tregs (in tissues and tumors) express high levels of immune checkpoint molecules OX-40, 4-1BB, GITR, CD30, TIGIT, ICOS and transcription factors such as PRDM1 (blimp-1) and BATF.^22,25–27,31–34^. Effector Tregs can mirror effector T helper (Th) cell polarization, by acquiring coexpression of FOXP3 with chemokine receptors and transcription factors associated with Th1 (CXCR3, T-bet), Th2 (GATA3, IRF4) or Th17 (RORC, STAT3) differentiation.^22,35–38^. This specific polarization has been associated with an enhanced suppressive efficacy towards the matching T effector response.^31,35–37,39–45^ Since most of these insights have been generated in mice, it is still largely unknown whether these principles also apply to human tissue Tregs. As recently highlighted by Sharma *et al.*,^46^ one of the most interesting, but yet elusive tissue sites for Treg function in humans is the maternal-fetal interface.

Pregnancy is a mystifying biological process when viewed from an immunological perspective, as it poses a unique challenge to the maternal immune system.^47,48^ While peripheral immunity against pathogens needs to remain intact,^49^ the semi-allogeneic fetus and placenta, which may harbor foreign paternal antigens, have to be tolerized. This suggests that the maternal immune response is delicately balanced during pregnancy, requiring tight regulation especially of the local immune response at the maternal-fetal interface, while maintaining systemic immune responses.^47,48,50^ The requirement for local regulation of the maternal immune response is underlined by the fact that human decidual T cells can recognize and actively respond to fetal cord blood cells.^51^ Accordingly, maternal Tregs have been shown to be indispensable for successful embryo implantation and pregnancy outcome in murine pregnancy, as they contribute to maternal-fetal tolerance on multiple levels.^47,52,53^ Specifically, depletion of maternal Tregs caused pregnancy loss due to immunological rejection of the fetus.^53,54^ In humans, maternal Tregs have been shown to be abundantly present in the gravid uterus,^55–62^ and normal human pregnancy is characterized by increased numbers of Tregs in the periphery and at the maternal–fetal interface.^56,61,63,64^ In patients with preeclampsia, a severe hypertensive disorder of pregnancy, and patients with recurrent miscarriages, Treg numbers are reduced both at the maternal-fetal interface and in the periphery,^57,65–70^ which implies that also in humans local presence of Tregs in the pregnant uterus is required for successful pregnancy outcome.

Previous studies investigating the maternal, uterine immune system in humans have been limited by the practical challenge of acquiring biopsy material of the uterine wall and have made use of the thin superficial decidual layer attached to the delivered placenta, which is heavily contaminated by fetal immune cells and may not be representative of the maternal Treg status during pregnancy. Moreover, the functional and transcriptomic profile of human Tregs from the maternal-fetal interface and its relation to Tregs from other human tissues remains to be elucidated. We hypothesize that resident Tregs in the gravid human uterus acquire a specialized functional profile to effectively regulate the local maternal immune response. Here, we investigated functional adaptation and specialization of highly purified human, exclusively maternal, resident uterine Tregs in myometrial biopsies from the maternal-fetal interface. We performed transcriptomic profiling and functional *in vitro* assays, as well as flow cytometry to study their phenotypic heterogeneity on protein level in single cell resolution. Furthermore, to identify tissue (site)-specific functional adaptation, we compared these Tregs to uterine Tregs from a distant uterine control site and maternal peripheral blood Tregs, in addition to the tissue- and site-matched resident CD4^+^ non-Treg T cells. Lastly, we compared the specific profile of functional adaptation of uTregs to known Treg signatures from other human and murine tissue sites, including tumors. In short, we identified a functional profile representing late-stage effector Treg differentiation, chronic activation, and Th1-like polarization, in uterine Tregs from the maternal-fetal interface, which revealed a remarkable overlap with tumor-infiltrating Treg signatures. Moreover, this functional adaptation represented not only local adaptation to the uterine tissue environment, but was specifically pronounced at the maternal-fetal interface, implying tissue site-specific adaptation within the pregnant human uterus.

## RESULTS

### Uterine Tregs are bona fide suppressive Tregs

The frequency of CD25^hi^FOXP3^+^ regulatory T cells (Tregs) within the CD4^+^ T cell population was similar between blood and uterine tissue and ranged from 2.5 to 13.5% (supplementary figure 1A). For transcriptomic analysis, the CD3^+^CD4^+^CD25^hi^CD127^−^ population (Tregs) and CD3^+^CD4^+^CD25^−^CD45RA^−^ memory T cells (Tconv) were FACS sorted from peripheral blood and myometrial biopsies from 5 women with uncomplicated pregnancies undergoing Caesarean section. In myometrium, Tconv were selected for CD69 positivity. The sorting strategy is shown in supplementary figure 1B. Confirming the maternal origin of the sorted cells, the female-specific gene XIST was highly expressed in all samples, whereas transcripts of the male-specific gene SRY were undetectable in all samples, including pregnancies with male offspring (supplementary figure 1C). Principal component analysis (PCA) of transcriptomic profiles showed that uterine Tregs (uTregs) from the maternal-fetal interface are clearly distinct from blood-derived Tregs (bTregs), and that also uterine T conv (uTconv) and blood-derived Tconv (bTconv) clearly cluster apart (figure 1A). Notably, PC1, mounting the difference between the cell sources, accounted for >60% of the variance, whereas PC2, explaining variance between Treg and Tconv populations, accounted for only 11% of the variance. To assess whether the sorted population of uTregs were bona fide Tregs, we analyzed enrichment of a published core Treg gene signature^78^ in uTregs compared to uTconv and bTregs by gene set enrichment analysis (GSEA). Expression of Treg core signature genes was not only enriched compared to uTconv, but, remarkably, also more pronounced in uTregs than in bTregs, indicating that uTregs are bona fide Tregs with enhanced expression of Treg core signature genes (figure 1B-C). Indeed, expression of many Treg markers from the published Treg signature^78^ was higher in uTreg than bTreg (figure 1D). Expression of the Treg-identifying molecules FOXP3 and CTLA-4 was confirmed to be higher in uTregs than bTregs on protein level (figure 1E-G). Also TIGIT, a key checkpoint molecule associated with specialized suppressive function,^79^ was highly expressed in uTregs, with the majority uTregs being positive for TIGIT (figure 1H). Consistently, GSEA showed significant enrichment of a previously identified TIGIT^+^ Treg signature (figure 1I).^79^ Suppression assays, although technically challenging due to low cell numbers, confirmed the suppressive potential of uTregs on proliferation and cytokine production of healthy donor peripheral blood-derived CD4^+^ T cells (Figure 1J and supplementary figure 1D-E). 2 out of 4 uTreg donors showed particularly high suppressive capacity of uTregs on cytokine production of IL-2, IL-10, IFNγ and TNFα, already at a 1:8 (Treg:Tconv) ratio, compared to bTregs. These results confirm that the sorted uTregs are bona fide functional Tregs, with enhanced expression of Treg signature genes.

**Figure 1.**
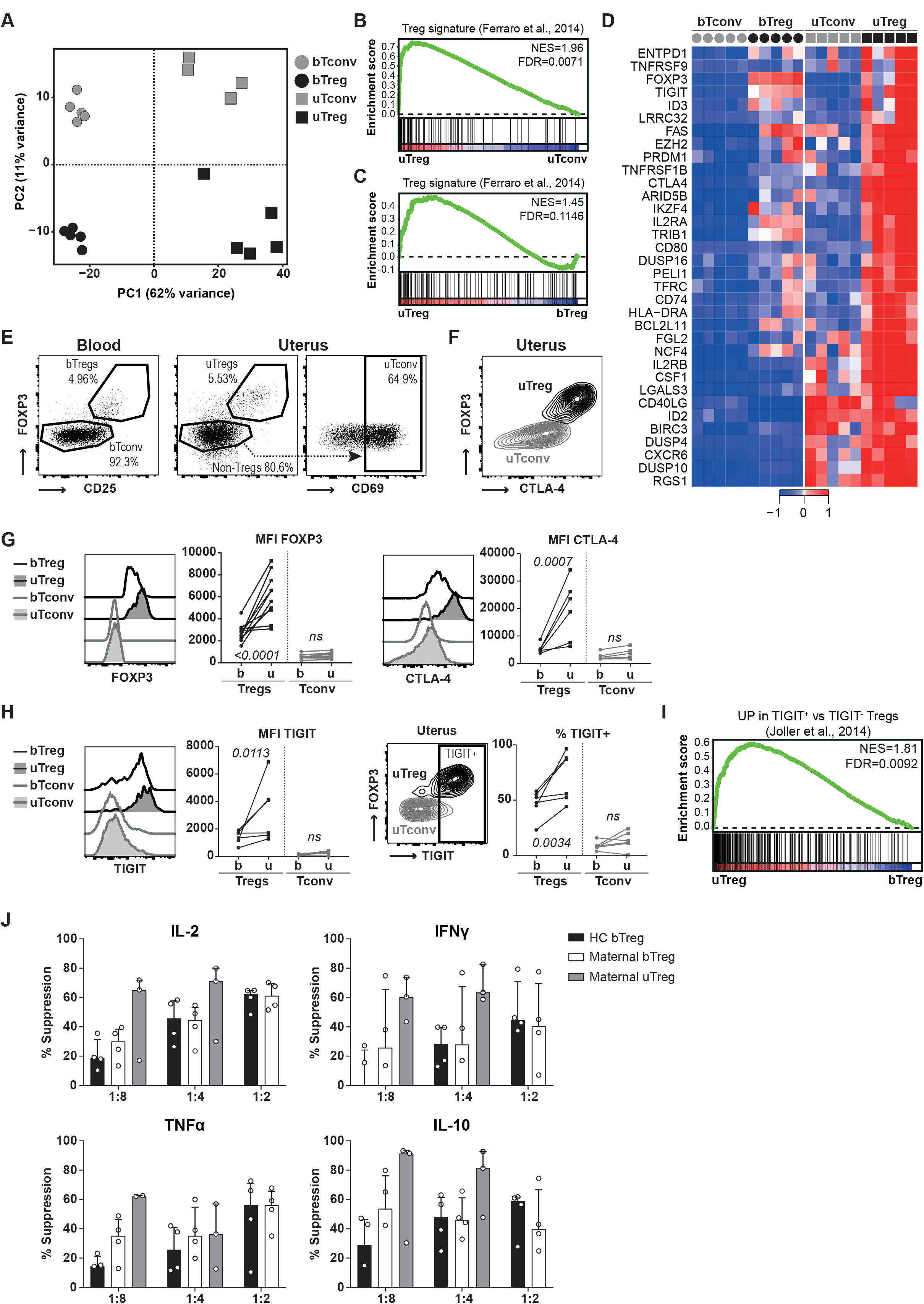
Tregs at the maternal-fetal interface are bona fide Tregs. **(A)** Principal component analysis of bTregs, bTconv, uTregs and uTconv. **(B+C)** Gene set enrichment analysis (GSEA) with published Treg signature gene set ^78^ comparing uTreg and uTconv **(B)** and uTreg and bTreg **(C)**. NES = normalized enrichment score. **(D)** Heatmap of genes in leading edge of GSEA analysis comparing enrichment of published Treg signature genes in uTregs and bTregs. Expression values were mean-centered and scaled per gene. **(E)** Gating strategy of bTregs, uTregs and uTconv. **(F)** Expression of CTLA-4 in uTregs. **(G)** Ex vivo protein expression of core Treg molecules FOXP3, CTLA4, and CD25 measured by flow cytometry. **(H)** Ex vivo protein expression of Treg signature molecule TIGIT measured by flow cytometry. **(I)** GSEA of TIGIT^+^ Treg signature.^79^ **(J)** Suppression assay assessing cytokine production of anti-CD3 stimulated (or unstimulated) healthy CD4^+^ T cells in the supernatant by multiplex immunoassay after 4 days of coculture with healthy donor bTregs, maternal bTregs, or uTregs at a 1:8, 1:4 and 1:2 ratio. *MFI = median fluorescent intensity*.

### The uTreg signature is indicative of an activated and effector Treg profile

To investigate the functional adaptation of uTregs to the specific environment of the maternal-fetal interface, we determined both their functional differentiation and (T helper) polarization, both of which may be influenced by the tissue environment.^13,17,18,33,80–82^ To identify the uTreg-specific transcriptional signature, we assessed their differential gene expression with both bTregs and uTconv. A large number of genes were differentially expressed between uTregs and bTregs (figure 2A). Differential gene expression analysis showed significant upregulation of 1966 genes and downregulation of 1997 genes in uTregs compared to bTregs (padj<0.05 and |Log2FC|>0.05). To isolate the uTreg specific signature, we also compared gene expression between uTreg and uTconv, yielding 465 upregulated genes, including the Treg-identifying genes FOXP3, IL2RA, CTLA4, TIGIT and IKZF2, and 103 downregulated genes in uTreg compared to uTconv (padj<0.05 and |Log2FC|>0.5; figure 2B) To pinpoint uTreg-specific genes, we overlapped their differentially expressed genes with bTreg and uTconv (figure 2C). This resulted in 236 genes specifically upregulated (225 after removal of duplicate genes) and 23 genes specifically downregulated in uTreg compared to both bTreg and uTconv (supplementary table 4). Among the 23 downregulated genes were ITGA6, IL7R, CCR7, TTC39C, PLAC8, ATF7IP2, ABLIM1, MGAT4A, PRKCB, GIMAPs as well as transcription factors TCF7, LEF1, and SATB1, indicating late-stage differentiation of Tregs.^83,84^ Pathway analysis of the 225 upregulated genes yielded cytokine signaling, TNF receptor signaling, and glycolysis as important upregulated pathways (figure 2D). Selected genes from the top 5 pathways included molecules related to Treg activation and effector differentiation, such as immune checkpoints of the TNF receptor superfamily (TNFRSF13B (TACI), TNFRSF18 (GITR), TNFRSF1B (TNFR2), TNFRSF4 (OX-40), TNFRSF8 (CD30), TNFRSF9 (4-1BB)) and HLA-DR, CD80, and LRRC32. Also molecules associated with suppressive capacity (CTLA4, ENTPD1, HAVCR2, IL10, IL2RA, LAG3, LAYN, LGALS1, PDCD1,and TOX2) were highly expressed in uTregs (figure 2E).^22,31^ Furthermore, cytokine receptors of the IL-1 and IL-2 family (IL1R1, IL1R2, IL1RAP, IL1RN, IL2RA, IL2RB) and specific chemokine receptors (CCR1, CXCR6) showed increased and specific expression in uTregs (figure 2E). Transcription factors that were specifically upregulated in uTregs included BATF, CEBPB, ETS2, ETV7, HES1, IKZF4, MAF, NFIL3, PRDM1, VDR, and ZBTB32 among others (figure 2E). This transcriptomic profile, and especially high expression of BATF, PRDM1, and immune checkpoint molecules, reflects previously identified crucial signatures of effector Treg differentiation and function, especially in tissues.^29,32,33,85–87^ We confirmed upregulation of the important immune checkpoints associated with effector Treg differentiation/chronic stimulation GITR, OX-40, 4-1BB, and PD-1, HLA-DR, and ICOS in uTregs on protein level, which again showed their specifically high levels in uTregs even compared to uTconv (figure 2F). Since increased expression of many of these genes pointed towards an activated phenotype, we confirmed this by demonstrating significant enrichment of published gene sets generated by activating Tregs *in vitro* with TCR stimulation or cytokine stimulation, in uTregs (supplementary figure 2, supplementary table 3).^88–92^ Taken together, these findings indicate that uTreg at the maternal-fetal interface have a highly differentiated transcriptional signature suggestive of a specialized function with high suppressive capacity and high responsiveness to environmental cues, which is reflective of late-stage effector differentiation and chronic activation.

**Figure 2.**
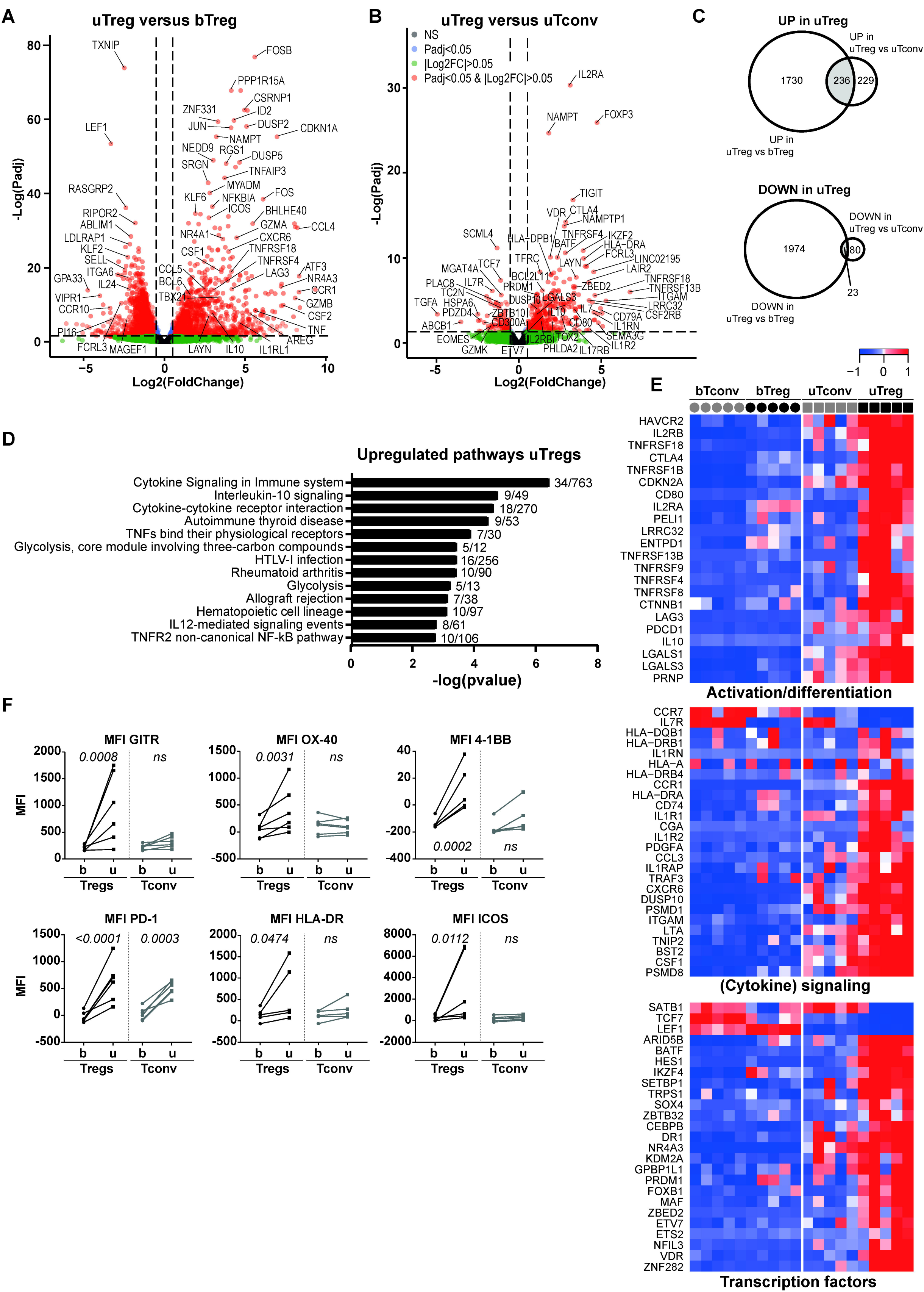
The uTreg core signature. **(A+B)** Volcanoplot of differential gene expression between uTregs and bTregs **(A)** or uTregs and uTconv **(B) (C)** Venn diagrams yielding genes specifically upregulated (padj<0.05 and Log2FC>0.05, upper panel) or downregulated (padj<0.05 and Log2FC<−0.5, lower panel) in uTreg compared to bTreg and uTconv. **(D)** Pathway analysis (ToppGene pathways) of 236 genes specifically upregulated in uTregs. P-values<0.05 after Bonferroni correction were considered significant. **(E)** Heatmap showing gene expression of genes in top 5 pathways and selected downregulated genes in the uTreg core signature, related to Treg activation or effector differentiation (upper panel), (cytokine) signaling (middle panel; including downregulated CCR7 and IL7R) and transcription factors (lower panel). Expression values were mean-centered and scaled per gene. **(F)**. Protein expression of GITR (TNFRSF18), OX-40 (TNFRSF4), 4-1BB (TNFRSF9), PD-1 (PDCD1), HLA-DR and ICOS. Padj of Two-way ANOVA with Tukey post hoc test. *MFI = median fluorescent intensity; NS = Not significant*.

### uTregs have a tissue-resident phenotype and share transcriptional specialization with uTconv

To examine whether uTregs at the maternal-fetal interface represent a resident population or rather transiently infiltrating cells, we assessed the expression of tissue-residency related markers and gene signatures. uTregs had a significantly higher gene and protein expression of key residency molecule CD69 than bTregs and bTconv, similar to uTconv (figure 3A-B). Expression analysis and GSEA with published human TRM signatures showed a pattern of upregulated and downregulated genes as previously described in CD4^+^ (and CD8^+^) TRM from the lung and skin (figure 3B-C),^4,15,93^ confirming the tissue-resident profile in uTregs as compared to bTregs. To further assess the tissue-specific adaptation of T cells at the maternal-fetal interface, we overlapped the genes that were significantly upregulated or downregulated in uTregs and uTconv compared to their counterparts from blood. The Venn diagrams in figure 3D show that a large proportion of upregulated and downregulated genes was shared between uTregs and uTconv (1032 up and 1348 down; padj<0.05 and |Log2FC|>0.05), which suggests that the specific tissue environment at the maternal-interface accounts for a significant part of their adapted transcriptional profile. Pathway analysis demonstrated that shared upregulated genes were involved in cytokine signaling (figure 3E). Downregulated pathways were reflective of ribosomal processes involved in RNA translation (supplementary figure 3). Taken together, uTregs have a TRM signature which reflects a shared adaptation to the tissue environment of the maternal-fetal interface between uTregs and uTconv.

**Figure 3.**
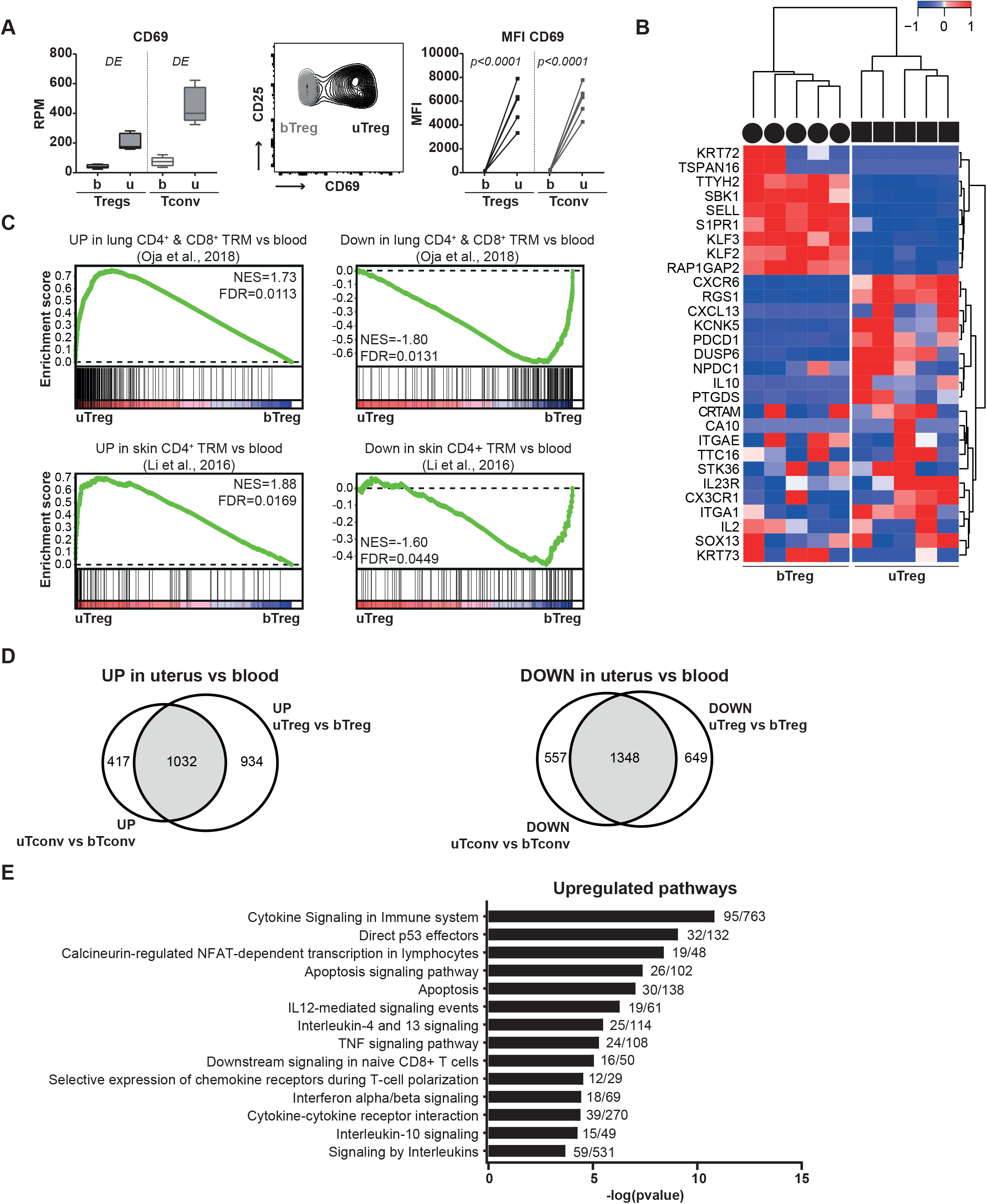
Tregs at the maternal-fetal interface have a tissue-resident profile. **(A)** Gene and protein expression of CD69 in sorted T cell populations. DE = differentially expressed padj<0.05. MFI = median fluorescent intensity. Two-way ANOVA with Tukey post hoc test. **(B)** Heatmap of published human core tissue-resident gene expression^4^ in uTreg compared to bTreg. Expression values were mean-centered and scaled per gene. **(C)** Gene set enrichment analysis (GSEA) with published genes identifying human lung CD4^+^ and CD8^+^ TRM compared to blood memory cells; left panel^93^) and genes upregulated in skin CD4+ TRM compared to blood CD4+ T cells (right panel^15^), in uTregs vs bTregs. NES = normalized enrichment score. **(D)** Venn diagrams of upregulated and downregulated genes (padj<0.05 and |Log2FC|>0.05) shared between Tregs and CD4^+^ Tconv from the maternal-fetal interface, compared to their blood-derived counterparts. **(E)** Pathway analysis (ToppGene pathways) of the 1032 shared upregulated genes in uTreg and uTconv. P-values<0.05 after Bonferroni correction were considered significant.

### uTregs mirror uTconv Th1 polarization with a predominance of T-bet+CXCR3+ Tregs

Effector Tregs can acquire different T helper phenotypes with coexpression of FOXP3 and lineage-defining transcription factors T-bet (TBX21, Th1), GATA3 (Th2), RORγt (RORC, Th17), as well as lineage-associated cytokine and chemokine receptors.^38^ We investigated whether uTregs and uTconv underwent a, possibly shared, T helper polarization. uTregs showed significantly increased expression of Th1-related TBX21 compared to bTreg, which mirrored the increased expression of TBX21 in uTconv compared to bTconv (figure 4A). Th2-related GATA3 and Th17-related RORC were not significantly differentially expressed between uTreg and bTreg (and uTconv and bTconv), although RORC showed a trend towards downregulation, which was confirmed on protein level (Figure 4A-B). Increased expression of T-bet was also confirmed on protein level, with 6-87% (median 22%) of uTregs showing positivity for T-bet (figure 4C-D). Also the Th1-related cytokine receptor IL18R1 was increased in both uTregs and uTconv compared to their counterparts in blood on gene and protein level (figure 4E). Investigation of chemokine receptor expression, which is related to both T helper polarization and tissue-specific homing,^94,95^ showed that chemokine receptors related to naive Tregs and lymphoid tissue environments CCR7 and CXCR5 were downregulated in uTregs compared to bTregs, on gene and protein level (figure 4F-G). Chemokine receptors upregulated in uTregs included CCR2, CCR5, CXCR3, CXCR4, CCR1, and CXCR6 (figure 4F and H), which largely mirrored expression by uTconv. CCR1 and CXCR6 were particularly upregulated in uTregs, both previously identified as part of the conserved murine tissue Treg signature.^21^ The Th1-associated CXCR3^35,96^ and Th1/inflammation-associated CCR5,^96–99^ had significantly higher gene and protein expression in uTregs and uTconv compared to their counterparts from blood (figure 4F and H). Although the variable percentage of T-bet^+^ Tregs suggests heterogeneity in uTreg subspecialization, virtually all uTregs (and uTconv) were positive for CXCR3 (84-100%, median 93%), and the majority expressed CCR5 (22-83%, median 62%) (figure 4I). Consistent with these findings, a previously published gene signature of T-bet^+^CXCR3^+^ Tregs from the pancreas of prediabetic mice was highly enriched in uTregs compared to bTregs (figure 4J).^37^ In conclusion, uTregs at the maternal-fetal interface show Th1 polarization mirroring uTconv, with high expression of Th1-related markers T-bet and CXCR3. Furthermore, uTregs express an array of chemokine receptors, some of which uTreg-specific and others shared with uTconv, with which they can integrate a variety of locally produced signals. uTreg and uTconv cells may therefore rely on both unique and shared cues to guide their migration to and retention at the uterine maternal-fetal interface.

**Figure 4.**
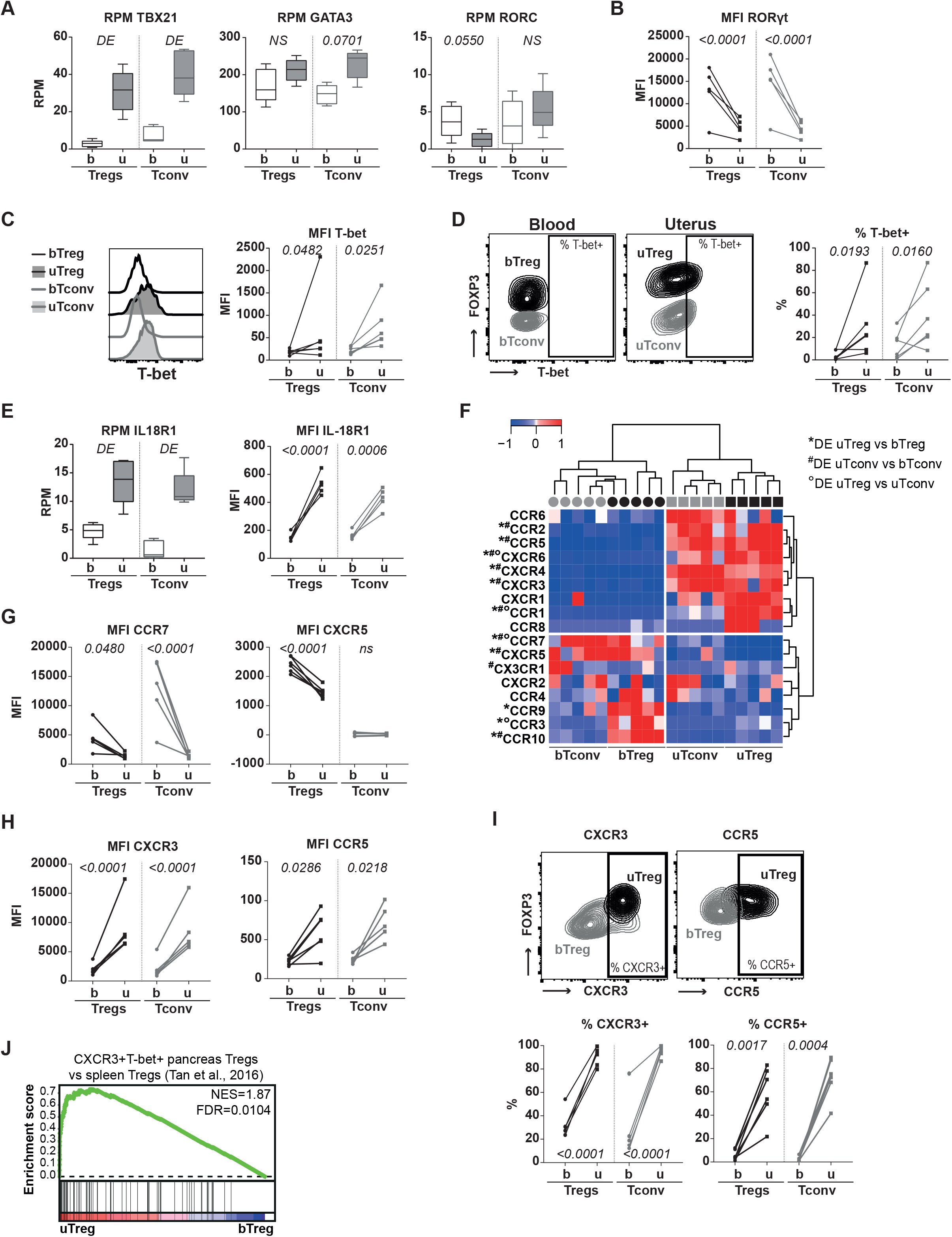
uTreg and uTconv polarization at the maternal-fetal interface. **(A)** Gene expression of lineage-defining transcription factors TBX21 (T-bet), GATA3 (GATA-3), and RORC (RORγt). P values from differential gene expression analysis. DE = differentially expressed padj<0.05. **(B-D)** Protein expression of RORγt **(B)** and T-bet **(C+D)**. MFI = median fluorescent intensity. P values of two-way ANOVA with Tukey posthoc test. **(E)** Gene and protein expression of IL18R1 (IL-18R1). **(F)** Heatmap showing gene expression of chemokine receptors. Expression values were mean-centered and scaled per gene. DE = differentially expressed padj<0.05. **(G-I)** Protein expression of chemokine receptors downregulated **(G)** and upregulated **(H+I)** in uTregs. P values of two-way ANOVA with Tukey posthoc test. **(J)** Gene set enrichment analysis with published gene set of CXCR3+T-bet+ Tregs from the pancreas of prediabetic mice,^37^ comparing uTregs and bTregs. NES = normalized enrichment score.

### The core uTreg signature from the maternal-fetal interface overlaps with tumor-infiltrating Treg signatures

uTregs at the maternal-fetal interface displayed profiles of effector Treg differentiation/chronic activation and Th1 polarization. We wondered whether these highly differentiated uTregs from the maternal-fetal interface would resemble Tregs from other human and murine tissue sites or would show a uniquely adapted profile. Well-studied murine tissue Treg populations include Tregs from visceral adipose tissue (VAT), muscle, and intestines.^12,13,21,80,100^ Each population has been shown to display a tissue-specific phenotype with expression of certain 1) transcription factors, 2) chemokine receptors and 3) preference towards a T helper (Th) lineage differentiation when compared to spleen Tregs.^12,13,17,21,80^ Next to these tissue-specific signatures, a murine PAN-tissue signature, shared by VAT, muscle and intestinal Tregs, was identified.^21^ GSEA in figure 5A shows that the shared murine PAN-tissue Treg signature was also strongly enriched in uTregs, again highlighting its generalized expression in tissue Tregs, apparently even conserved across species. Overlaying significantly upregulated genes in uTreg (versus bTreg) with the murine tissue-specific or tissue-shared Treg signatures,^21^ yielded a large amount of shared genes between uTregs and murine VAT-, colon- and muscle-derived Tregs (figure 5B, numbers in each field represent overlap of the specific field with significantly upregulated genes in uTreg). 59 genes were shared among all 3 murine tissues and uTregs, including IL1RL1 (receptor for IL-33, ST2), AREG, IL10, IRF4, GZMB, TNFRSF9, BHLHE40, NR4A1, NR4A3, and CCR2, many of which have been described as crucial regulators for effector and/or tissue Treg function (figure 5C).^13,32,40,85–87,101^ 12 of the 59 genes were even part of the uTreg-specific core signature as defined in figure 2 (CCR1, CXCR6, ELL2, FGL2, GEM, IL10, LAPTM4B, SNX9, TNFRSF8, NFIL3, NR4A3, and PRDM1). This indicates that uTreg display features of tissue adaptation, which are highly conserved across tissues and species.

**Figure 5.**
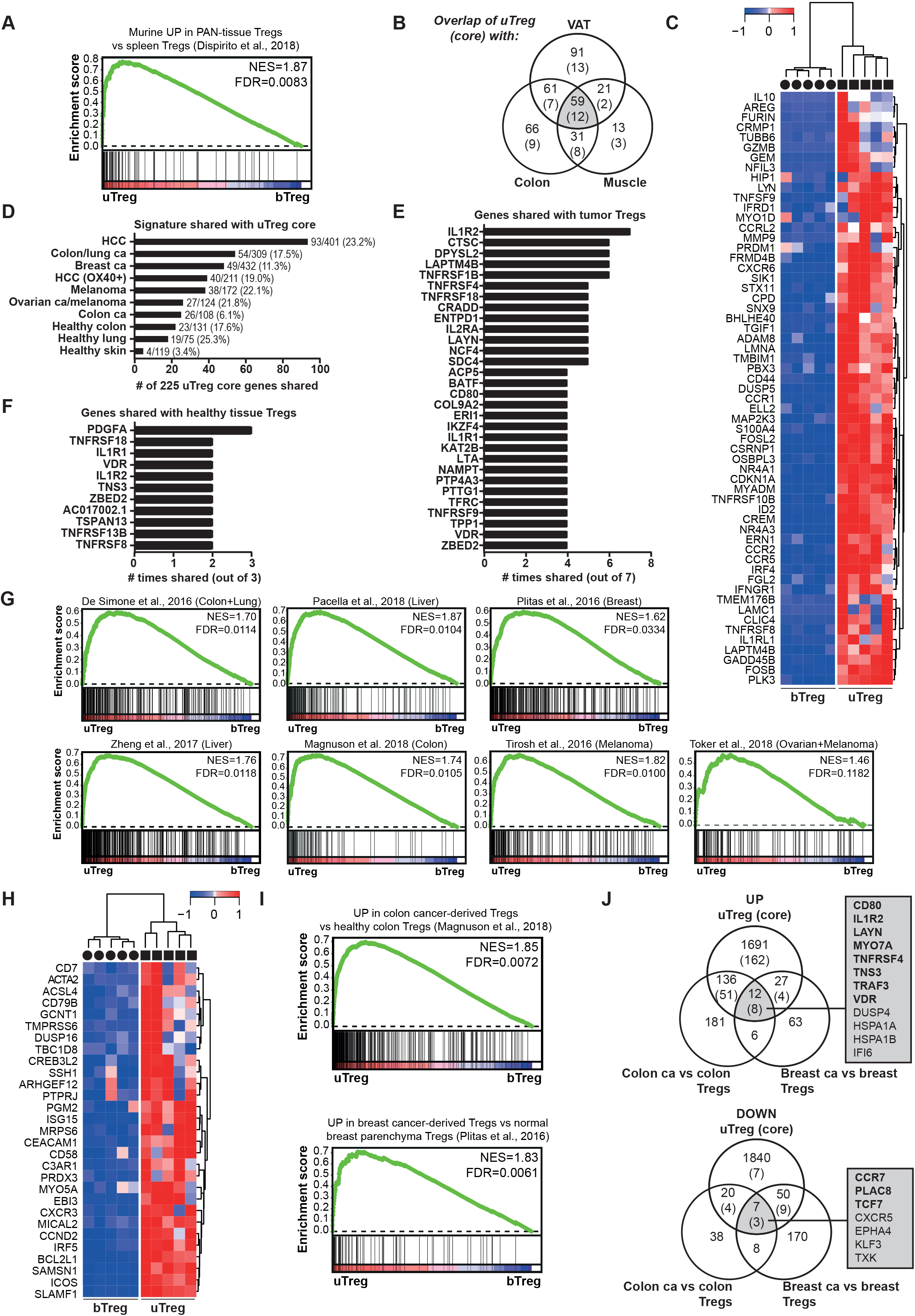
uTregs have a functional profile similar to tumor-infiltrating Tregs. **(A)** Gene set enrichment analysis with a published murine PAN-tissue gene signature,^21^ comparing uTregs and bTregs. **(B)** Venn diagram showing the numbers of genes upregulated in uTregs compared to bTregs (padj<0.05) (and genes in the uTreg core signature in parentheses), which are represented in tissue-specific and tissue-shared published murine gene signatures.^21^ VAT = Visceral adipose tissue. **(C)** Heatmap showing the expression of the 59 genes that were part of the murine PAN-tissue signature and upregulated in uTregs compared to bTregs (padj<0.05).^21^ Expression values were mean-centered and scaled per gene. **(D)** The number of genes shared between the uTreg core signature and published human TITR signatures or healthy tissue Treg signatures. ^14,15,25,102–107^ Numbers behind bars indicate the number of shared genes out of the total number of genes in the specific signature. **(E)** The genes that were most often shared between the uTreg core signature and human TITR signatures (shared in >3/7 signatures). **(E)** The genes that were most often shared between the uTreg core signature and human healthy tissue Treg signatures (shared in >1/3 signatures). **(G)** Gene set enrichment analysis (GSEA) with published TITR-specific signatures in uTregs vs bTregs.^25,102–107^ NES = normalized enrichment score. **(H)** Heatmap showing expression of genes in the leading edge of >2/7 GSEA analyses from **(G)**, which were not represented in the uTreg core signature. Expression values were mean-centered and scaled per gene. **(I)** GSEA with published gene signatures specific to Tregs from tumor-tissue compared to the healthy tissue counterpart in uTregs vs bTregs.^26,107^ **(J)** Venn diagrams showing shared genes between uTregs and genes specifically upregulated in Tregs from tumor-tissue compared to the healthy tissue counterpart.^26,107^

In the human setting, the investigation of tissue-derived Tregs, especially from healthy tissues, has proven challenging, and only limited data are available. To assess how the uTreg tissue profile compared to other human Treg tissue profiles, we analyzed enrichment of the three previously published gene sets of significantly upregulated genes in skin, colon and lung Treg compared to blood Treg (supplementary table 3).^14,15^ All three of these signatures were significantly enriched in uTregs compared to bTregs, indicating that the tissue profile of uTregs shows similarities with human Tregs from various tissue sites (supplementary figure 4). Human Tregs infiltrating the unique tissue-environment of tumors (TITR) have been studied slightly more extensively. Comparison of genes significantly upregulated in uTregs versus bTregs with seven recently published gene signatures of TITR infiltrating a variety of human tumors (supplementary table 3),^25,102–107^ yielded a remarkable overlap with each of the TITR signatures with up to 65% of genes shared with uTregs (supplementary table 5). Of the 41 genes that were shared among ≥4 of the 7 TITR signatures (supplementary table 6), a notable 31 were also part of the 225 genes in the uTreg core signature. Figure 5D shows the genes that were shared between the uTreg core signature and each of the TITR signatures and healthy tissue-derived Treg signatures. Remarkably, 93 (41.3%) of the 225 core uTreg genes were overlapping with specifically upregulated genes from HCC-infiltrating Tregs,^105^ 54 with the unique TITR signature identified by De Simone et al,^25^ 49 with breast cancer TITR genes,^102^ and 40 with OX-40^+^ Treg from cirrhotic/tumor liver tissue (figure 5D).^106^ Importantly, the 225 uTreg core signature genes showed less overlap with healthy tissue-derived Treg specific signatures from human healthy colon, lung and skin. The genes that were most often shared between uTregs and TITR were IL1R2 (7/7), TNFRSF1B, CTSC, DPYSL2, LAPTM4B (6/7), TNFRSF4, TNFRSF18, LAYN, IL2RA, ENTPD1, NCF4, SDC4 and CRADD (5/7) (figure 5E), whereas with healthy tissue-Treg signatures PDGFA was most often shared (3/3) (figure 5F). GSEA showed that also many of the non-overlapping genes from the published TITR signatures were significantly enriched in uTreg compared to bTreg (figure 5G). Genes in the leading edge of ≥3/7 tumor-specific GSEA analyses that were highly expressed in uTregs compared to bTregs, but not part of the uTreg core (mostly because their high expression was shared with uTconv), are shown in figure 5H. These included CREB3L2 (6/7) EBI3, GCNT1, ICOS (5/7), ACTA2, ARHGEF12, BCL2L1, CCND2, PRDX3, SLAMF1 (4/7), CXCR3, CD7, CAECAM1, CD79B, and MICAL2 (3/7), amongst others. Remarkably, genes specifically upregulated in breast cancer-infiltrating Tregs compared to Tregs from normal breast parenchyma or significantly upregulated in colon cancer Tregs compared to healthy colon Tregs showed a particularly high enrichment in uTregs, suggesting that uTregs are not just similar to Tregs from breast or colon tissue, but specifically to the highly differentiated/activated Tregs from the tumor environment (figure 5I).^26,107^ By overlapping these cancer-versus-healthy tissue Treg signatures with significantly upregulated genes in uTregs (versus bTregs), we identified 12 ‘cancer-specific’ genes expressed by uTregs (figure 5J): CD80, IL1R2, LAYN, MYO7A, TNFRSF4, TNS3, TRAF3, VDR, DUSP4, HSPA1A, HSPA1B, and IFI6. The first 8 of these were also part of the uTreg-specific core signature, again highlighting the specificity c.q. importance of receptors IL1R2, LAYN, TNFRSF4, CD80 and transcription factor VDR for human Tregs in a tumor(-like) microenvironment. Also tumor-specific downregulated genes were shared with the uTreg core signature: CCR7, PLAC8, and TCF7. In conclusion, these results indicate that uTreg from the maternal-fetal interface have a transcriptional core signature which is shared specifically with the specialized transcriptional profile of tumor-infiltrating Tregs.

### Uterine Tregs show site-specific adaptation within the uterus

Next we wondered whether uTreg would be merely adapted to the microenvironment in uterine tissue, or specifically adapted to the tissue site at the maternal-interface. To investigate this site-specific adaption within one organ, we compared uTregs from the maternal-fetal interface, i.e. placental bed (^pb^uTregs), to uTregs from a distant uterine site, i.e. the incision site made during Caesarean section (^inc^uTregs). Confirmation of Treg identity and TRM signature for ^inc^uTregs is shown in supplementary figure 5A-F. The differentially expressed genes between ^inc^uTregs and bTregs were similar to those between ^pb^uTregs and bTregs (figure 6A). Also PCA showed that gene expression profiles of ^pb^uTregs and ^inc^uTregs were rather similar, compared to bTregs (figure 6B). However, direct comparison of ^pb^uTregs and ^inc^uTregs revealed a substantial difference between the two populations (figure 6B-C). First, protein expression of the core Treg transcription factor FOXP3 was lower in ^inc^uTregs than ^pb^uTregs, comparable to bTegs (figure 6D). This was not due to ^inc^uTreg contamination with bTregs, since expression of CD69 was similar between ^pb^uTregs and ^inc^uTregs (supplementary figure 5D). Protein expression of other core Treg genes CTLA4 and TIGIT was also lower in ^inc^uTregs than ^pb^uTregs (figure 6D). This indicates that ^pb^uTregs, derived from the maternal-fetal interface, have a more pronounced expression of Treg signature markers, suggesting enhanced activation/differentiation in comparison with their uterine counterparts from the incision site. Differential gene expression analysis revealed 558 upregulated and 125 downregulated genes in ^pb^uTregs versus ^inc^uTregs (figure 6E). The heatmap in figure 6F shows a selection of previously highlighted genes in this manuscript that proved to be differentially expressed between ^pb^uTregs and ^inc^uTregs. These results suggest that Tregs cannot only adapt to the microenvironment within a certain tissue, but will specifically adapt to the environmental cues at a specific tissue site. Pathway analysis showed that upregulated genes ^pb^uTregs versus ^inc^uTregs were related to PD-1 signaling, cytokine signaling, TCR signaling, and T helper cell differentiation (supplementary figure 5G). Indeed, PD-1 was higher expressed in ^pb^uTregs and ^inc^uTregs on gene and protein level (figure 6F-G), and GSEA showed enrichment of a TCR-activated Treg signature in ^pb^uTregs compared to ^inc^uTregs (supplementary figure 5H). Furthermore, ^pb^uTreg-specific core genes associated with effector Treg differentiation including TNFRSF4 (OX-40 protein, figure 6H) and transcription factors BATF, MAF, PRDM1, and VDR, among others, were significantly higher expressed in ^pb^uTreg than in ^inc^uTreg (figure 6F), again suggesting that ^pb^uTreg show more pronounced differentiation towards an effector Treg phenotype. Since ^pb^uTregs appeared to be especially differentiated at the maternal-fetal interface, we assessed whether the TITR-like profile of ^pb^uTregs was also more pronounced than in ^inc^uTregs. Remarkably, 5 out of 7 tested published TITR signatures were significantly enriched in ^pb^uTregs compared to ^inc^uTregs (P<0.05; figure 6I). More specifically, GSEA with signatures differentiating between TITR and their counterparts from a matched healthy tissue site, showed significant enrichment in ^pb^uTregs compared to ^inc^uTregs (figure 6J). CCR8 and ICOS, which were present in in 6 out of 7 TITR signatures, as well as TNFRSF18 (GITR), were significantly higher expressed in ^pb^uTregs than in ^inc^uTregs and bTregs on protein level (figure 6K). CCR8 has been shown to be highly enriched in tumor Treg cells and associated with a poor prognosis in several cancers.^25,26,81^ Thus, ^pb^uTregs at the maternal-fetal interface specifically acquire a highly differentiated effector profile similar to tumor-infiltrating Tregs, which is more pronounced even compared to a uterine tissue site distant from the maternal-fetal interface.

**Figure 6.**
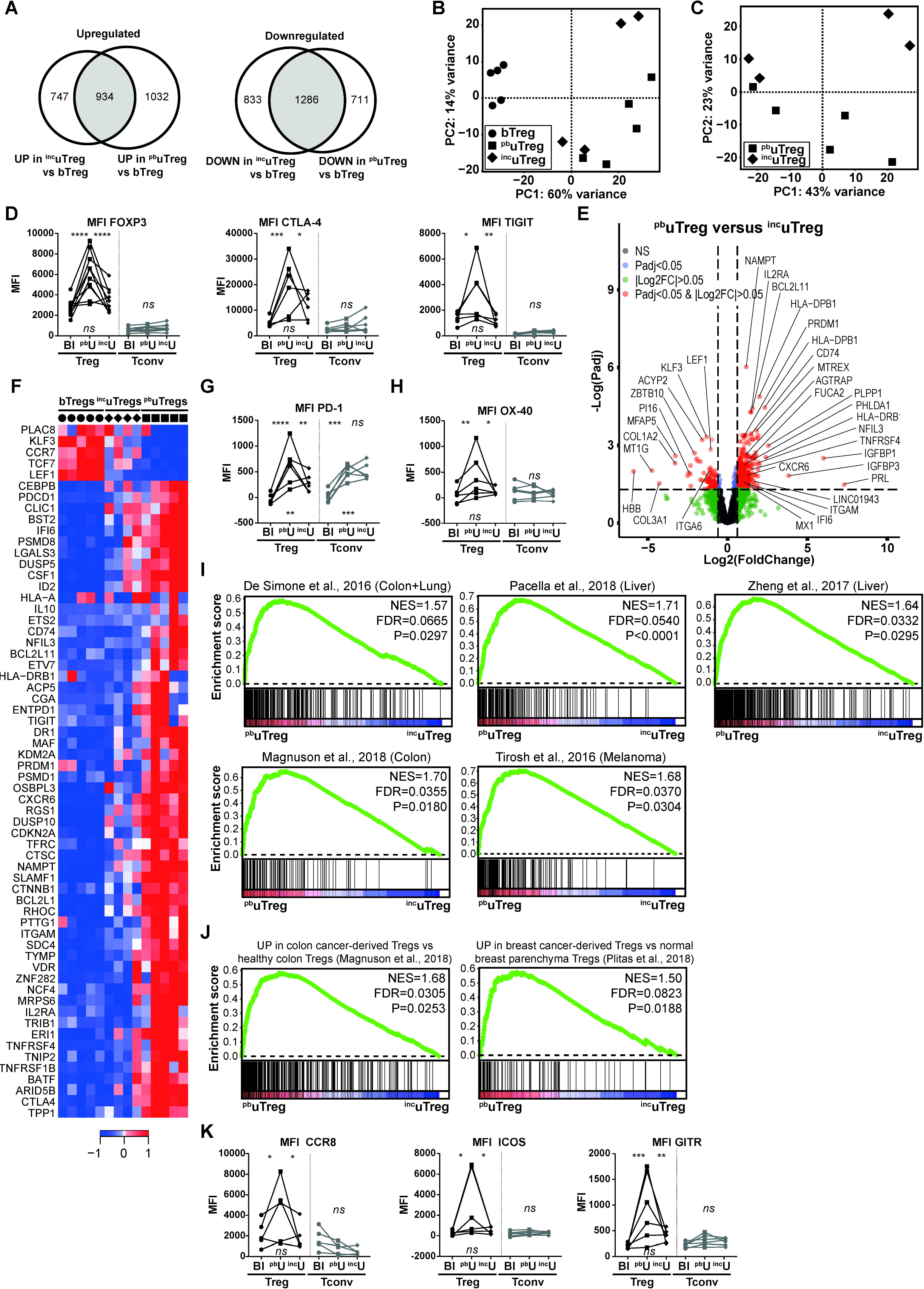
uTregs show site-specific adaptation to the maternal-fetal interface. **(A)** Venn diagrams of genes upregulated (left panel) and downregulated (right panel) in both ^inc^uTregs and ^pb^uTregs compared to bTregs. **(B)** PCA of bTregs, ^pb^uTregs and ^inc^uTregs. **(C)** PCA of ^pb^uTregs and ^inc^uTregs. **(D)** Protein expression of FOXP3, CTLA-4, and TIGIT. Padj of Two-way ANOVA with Tukey post hoc test for protein. *Left upper p-value: blood vs placental bed; right upper p-value: placental bed vs incision site; lower p-value: blood vs incision site. MFI = median fluorescent intensity; NS = Not significant*. **(E)** Volcanoplot of differentially expressed genes between ^pb^uTregs and ^inc^uTregs. **(F)** Heatmap with previously highlighted genes in this manuscript which were differentially expressed between ^pb^uTregs and ^inc^uTregs. Expression values were mean-centered and scaled per gene. **(G+H)** Protein expression of PD-1 **(G)** and OX-40 **(H)**. Padj of Two-way ANOVA with Tukey post hoc test for protein. *Left upper p-value: blood vs placental bed; right upper p-value: placental bed vs incision site; lower p-value: blood vs incision site. MFI = median fluorescent intensity; NS = Not significant*. **(I)** Gene set enrichment analysis (GSEA) with published TITR-specific signatures in ^pb^uTregs vs ^inc^uTregs. ^25,102–107^ NES = normalized enrichment score. **(J)** GSEA with published gene signatures specific to Tregs from tumor-tissue compared to the healthy tissue counterpart in ^pb^uTregs vs ^inc^uTregs.^26,107^ **(K)** Protein expression of CCR8, ICOS and GITR. Padj of Two-way ANOVA with Tukey post hoc test for protein. *Left upper p-value: blood vs placental bed; right upper p-value: placental bed vs incision site; lower p-value: blood vs incision site. MFI = median fluorescent intensity; NS = Not significant. ****P<0.0001, ***P<0.001, **P<0.01, *P<0.05*.

## DISCUSSION

Here, we demonstrate for the first time that, in pregnancy, human uterine Tregs have a highly differentiated transcriptional profile, which is specifically enriched at the maternal-fetal interface and is reminiscent of the specialized and highly effective profile of tumor-infiltrating Tregs. With these findings we answer a long-standing question on how Tregs are functionally specialized at the maternal-fetal interface to modulate local effector T cell responses, preventing an allo-reaction against the fetus. Moreover, we introduce the novel concept of site-specific adaptation of Tregs within one organ or tissue. This again substantiates the notion that Tregs are capable of adapting their transcriptional program driven by micro-environmental cues.^12,14,16,21,22^

We have demonstrated that uTregs at the maternal-fetal interface display a highly activated and late-stage differentiated effector profile (as suggested by expression of BATF and PRDM1, and downregulation of SATB1), ^32,34,83,84,86^ with increased expression of molecules associated with enhanced suppressive capacity (CTLA4, ENTPD1, HAVCR2, IL10, LGALS1, TIGIT) and abundant expression of TNFR superfamily members ((TNFRSF13B (TACI), TNFRSF18 (GITR), TNFRSF1B (TNFR2), TNFRSF4 (OX-40), TNFRSF8 (CD30), TNFRSF9 (4-1BB)). Also others have found that non-lymphoid-tissue Tregs display an activated phenotype compared to lymphoid-organ and circulating Tregs,^13,18,80^ and both BATF and the TNFRSF-NF-κB signaling axis have been described as crucial in the survival of Tregs and maintenance of a stable effector Treg phenotype, especially in tissues.^16,28,32,33,85,86,108^ It is now recognized that Tregs adapt to their tissue environments, with common adaptations across many tissues, such as increased expression of IL10, IL1RL1 (encoding ST2, an IL-33 receptor subunit), AREG (encoding amphiregulin), CTLA4, TIGIT, BATF and IRF4, and a low expression of LEF1 and TCF7 compared to lymphoid tissue Tregs, but, importantly, also tissue-specific signatures.^12,13,16,17,21^ These tissue-specific transcriptomic profiles counter the notion that tissue Tregs merely have a more activated, effector or memory state than lymphoid-organ Tregs. Rather, they have a specialized adapted program,^100^ likely matching the specific requirements of a certain tissue site.^10,17,21,109,110^

Although site-specific distribution of T cell composition and maturation in the human intestinal tract has been previously reported,^111^ to our knowledge, the concept of site-specific transcriptional adaptation of Tregs within one tissue or organ is novel, taking into account that tumors represent a completely altered tissue and not a different site within the same organ. We show that uTregs display features suggestive of a high responsiveness to micro-environmental cues, such as a range of TNF receptor superfamily members and chemokine receptors. With such a matrix of options to detect signals from the microenvironment, Tregs are likely able to adjust not only to the tissue or organ of their residence, but even to specific sites within that tissue, based on the cues provided by surrounding cells. Most likely, the implantation of the placenta, i.e. the multitude of signals produced by myometrium-invading trophoblast,^112^ are the primary cues effectuating micro-environmental changes at the maternal-fetal interface. It has been shown that trophoblast attracts Tregs to the maternal-fetal interface by production of hCG and CXCL16, the ligand for CXCR6.^113,114^ Moreover, *in vitro* co-culture of HLA-G^+^ extravillous trophoblast with CD4^+^ T cells increased Treg numbers and the FOXP3 expression level,^115,116^ indicating that Tregs may also be locally induced or expanded by trophoblast. Thus, it is likely that signals produced by invading trophoblast at the maternal-fetal interface account for at least some of the site-specific transcriptional adaptations in uTregs.

The T helper response at the maternal-fetal interface has been previously suggested to be skewed away from a pro-inflammatory Th1 response, to prevent a pathogenic alloreaction against the fetus, resulting in a Th2 dominant response during the second trimester. However, during the third trimester, a pro-inflammatory Th1 response was described to be essential for initiation of labor.(reviewed in^48^) In line with this, our findings indicate that the T helper response in the uterus at term is dominated by Th1 polarization, but that it is very well-controlled. The highly differentiated population of uTregs at the maternal-fetal interface appears to be specifically equipped to effectively suppress Th1 responses. Most importantly, although we observed heterogeneity of T-bet protein expression in uTreg, CXCR3 expression was remarkably homogeneous, with 84-100% of uTregs being CXCR3^+^. CXCR3 (and T-bet) expressing Tregs have been shown to be especially adept to suppress Th1 responses.^35,37,45,96^ Furthermore, the majority of uTregs expressed TIGIT, OX-40 and/or CCR5. Tregs expressing TIGIT have been described to preferentially inhibit Th1 and Th17 responses,^79^ a subpopulation of OX-40 expressing Tregs are thought to differentiate into Th1-suppressing Tregs,^117^ and also CCR5 expression on Tregs has been associated with more effective suppression of Th1 responses.^97^ Thus, the necessary pro-inflammatory Th1 response at the maternal-fetal interface at term appears to be controlled by specifically differentiated and Th1-polarized Tregs. So far, Th1-like Tregs have been described mainly in inflammatory environments, such as infections, autoimmune diseases and transplantation reactions,^45,96,118–120^ whereas tissue-resident Tregs have been mostly characterized as being Th2-skewed (VAT, muscle)^17,21^ or Th17 skewed (intestines).^80^ DiSpirito *et al.* however recently also identified a subset of T-bet expressing Tregs in muscle and colon,^21^ indicating that they can be present also in steady-state tissues.

We are the first to study exclusively maternal, myometrial tissue-resident Tregs from the maternal-fetal interface. Although Tregs at the human maternal-fetal interface have been studied previously, investigations had to resort to the use of more easily accessible decidua, due to the difficulty of acquiring human myometrium. Since decidual tissue is of fetal origin, it may not only be contaminated with fetal immune cells, but it also does not allow for studying the unique, and specifically maternal, uterine environment underlying the placenta, in which the complex process of spiral artery remodeling takes place. The only publications that we know of investigating FOXP3 expression in actual human placental bed biopsies demonstrated that the percentage of FOXP3^+^ T cells was significantly decreased in patients with pre-eclampsia, and FOXP3 mRNA expression was reduced in endometrial biopsies of infertile women, highlighting the importance of functional Tregs for a healthy pregnancy.^57,121^ From human decidual data, it is known that the frequency of clonally expanded populations of effector Treg cells is increased in decidua of 3^rd^ trimester cases compared to 1^st^ trimester cases.^122^ Decidual Tregs were found to display a more pronounced suppressive phenotype than in blood, with increased expression of FOXP3, CTLA-4, CD25, HLA-DR, ICOS, GITR, and OX-40, which recapitulates our findings.^58,59,63,123^ Very recently, three types of functional regulatory T cells were identified at the human maternal-fetal interface, of which the CD25^hi^FOXP3^+^ population matches the here studied population.^116^ Decidual CD25^hi^FOXP3^+^ Tregs effectively suppressed CD4^+^ and CD8^+^ T cell proliferation and IFN⍰ and TNF⍰ production. Transcripts identified by qPCR array as specific for this subset were IL2RA, FOXP3, TIGIT, CD39, LRRC32, ST2, BATF, and CCR8, as well as increased expression of CCR5, IL10, and GITR compared to blood Tregs,^116^ which confirms our findings of an activated Treg phenotype at the maternal-fetal interface. A previously published study investigating chemokine receptor expression of CXCR3, CCR4, and CCR6 in decidual Tregs by flow cytometry, showed that CCR6^−^CXCR3^+^ Th1 cells were increased, CCR6^+^CCR4^+^ Th17 cells were nearly absent, whereas CCR4^+^ Th2 frequencies were similar in blood and decidua,^58^ which is also in line with our findings. Remarkably, in murine gravid uterus, CCR5 expression on Tregs was related to suppressive capacity: CCR5^+^ effector Tregs were more suppressive compared to their CCR5^−^ counterparts.^124^ Furthermore, also in human TITR CCR5 was highly expressed, even specifically compared to the healthy colon-derived Tregs, and also here CCR5 expression on Tregs correlated with an increased suppressive capacity.^107,125^ Taken together, this indicates that the here identified activated phenotype of myometrial uTregs has overlapping characteristics with decidual Tregs.

We observed that uTregs from the maternal-fetal interface are highly responsive to their micro-environment. They display a peculiar differentiated effector phenotype similar to TITR, defined by high gene expression of IL1R2, LAYN, CD80, VDR, and TNFRSF4, amongst others, with specific enrichment of TITR signatures compared to Treg signatures from matched, unaffected tissue sites. This observation may be explained by recent insights on the similarity of the immune environment at the maternal-fetal interface and tumors.^48^ Both the receptivity of the myometrium towards implantation of the blastocyst and the invasiveness of the trophoblast show striking similarities with implantation of tumor metastases in healthy tissues.^126,127^ Tumor cells can modulate their immune environment into an anti-inflammatory milieu and have been shown to recruit and/or induce suppressor cells among which high numbers of suppressive Tregs.^128,129^ Just as in tumors, a tolerogenic mode of antigen presentation with indirect allorecognition of low levels of antigens predominates at the maternal-fetal interface.^130^ Also others have reported striking similarities between the early Treg responses to embryo and tumor implantation.^54^ And not only Tregs, but also neutrophils in decidua basalis have been shown to be similar to tumor-associated neutrophils.^131^ These findings imply that the micro-environment at the maternal-fetal interface may be a unique mammalian tissue site that under challenged, but physiological conditions resembles a tumor micro-environment. The tumor environment represents an actively remodeling tissue site distinct from a steady-state tissue, with low-grade inflammation, and newly infiltrating/invading cells. These dynamic characteristics are shared with the maternal-fetal interface and may account for the unique transcriptional adaptation of Tregs.

Although we observed global changes in gene expression patterns in uTregs, flow cytometry revealed an expression gradient of many markers across the uTreg population, suggesting that uTregs consist of a heterogenic population with different stages of differentiation and possibly sub-specialization. Single cell sequencing techniques and mass cytometry are indeed starting to reveal the heterogeneity of Treg populations in tissues and tumors.^16,27,104,105,132–134^ Treg heterogeneity may even contribute to their suppressive potential: IL-35 (encoded by EBI3) and IL-10 expressing heterogeneic Treg subpopulations in the tumor micro-environment have been recently shown to induce CD8^+^ Tconv exhaustion.^135^ Considering the striking similarities between uTreg and TITR and the high expression of IL-35 and IL-10 in uTregs, this could also be a potential mechanism employed by uTregs at the maternal-fetal interface, which is in line with a recent report showing partial CD8^+^ effector dysfunction in the human decidua.^136^

A unique strength of our study is that we were able to study a highly specific and pure maternal Treg cell subset isolated by FACS sorting from human tissue biopsies, of which we compared the transcriptomic profiles by state of the art low-input CEL-Seq protocol, not only to their counterpart in blood, but also to a tissue and site-specific Treg control population and matched Tconv. Moreover, we have validated our key findings on protein level in single cell resolution by flow cytometry. This revealed that some of the differences found on RNA level were even more pronounced on protein level. As we were only able to study T cells from biopsies acquired in term pregnancies, due to the practical limitation of delivery of the infant and placenta, it would be interesting to investigate term-dependent changes in uTreg profiles in future studies. It should be taken into account that protocols for tissue digestion may induce transcriptional changes.^137^ However, many of the uTreg specific genes identified here, were previously found not to be affected in human lung-derived Tregs by a tissue digestion protocol which was similar to but harsher than the isolation method used here.^14^

Our findings have important implications. TITR are currently under heavy investigation as targets in cancer immunotherapy. However, we demonstrate that signatures identified in TITR are not as unique as previously assumed, and that they may be shared by Tregs with specialized functions in other human tissues that may still be unknown. On the other hand, our results may lead to new targets for cancer immunotherapy, since profiling of Tregs in a variety of tissues under physiological, but not necessarily steady-state conditions, may help to identify truly TITR-specific expression patterns. Moreover, increased understanding of immunoregulatory mechanisms at the maternal-fetal interface during healthy pregnancy gives not only unique insights into human immunobiology of pregnancy, but also aids to elucidate the pathological changes in Tregs in pregnancy disorders such as preeclampsia, fetal growth restriction or recurrent miscarriage, as many studies have pointed towards a role for Treg defects of deficiency in these disorders.^65–67,113,122,138,139^ Lastly, functional adaptation of human Tregs to different tissues and specific tissue sites is still largely unexplored. The receptivity of Tregs to their environmental stimuli and subsequent sub-specialization may be exploited for therapeutic purposes. In conclusion, we have shown that human Tregs show functional adaptation with tumor-infiltrating-like features specifically at the maternal-fetal interface, which introduces the novel concept of tissue site-specific transcriptional adaptation of human Tregs.

## Acknowledgements

We thank Michal Mokry, Noortje van Dunen and Nico Lansu for their help with RNA sequencing. We thank the multiplex core facility, and especially Jeroen van Velzen and Pien van der Burght for their advice and (both practical and moral) support during FACS sorting. We thank Tatjana Vogelvang for her help in the collection of samples at the Diakonessenhuis in Utrecht and Arie Franx for his involvement in establishing the SPAR study initiative at the Wilhelmina Children’s Hospital.

We have no specific funding to report. LB recruited and included patients, and collected clinical data. JW, LB, RS and LvdB performed all wet-lab experiments. MM performed the RNA sequencing and helped with data-analysis. JW performed all data-analyses and wrote the manuscript. PN consulted on biopsy preparation, tissue integrity and uterine T cell distribution and phenotype. BvR and FvW supervised JW, LB, LvdB and RS, and were closely involved in setting up the study protocol, collection of data, data analysis and writing of the manuscript. All authors critically revised the manuscript. All authors declare no other support from any organization for the submitted work than the grants reported in the funding section; no financial relationships with any organizations that might have an interest in the submitted work in the previous three years, no other relationships or activities that could appear to have influenced the submitted work. The datasets generated for this study have been submitted to the Gene Expression Omnibus (accession code GSE…‥ (to be provided when submission is final)).

## MATERIALS AND METHODS

### Participants and biopsies

This study is part of the Spiral Artery Remodeling (SPAR) cohort study, which is an on-going effort to investigate the adaptation of the uterus to placental development by obtaining site-specific uterine biopsy samples in women undergoing Caesarean section. Detailed description of the study set-up and protocol was previously published.^71^ For this analysis, we included 20 women who delivered by elective Caesarean section, i.e. without any contractions or other signs of labour such as rupture of membranes, after an uneventful pregnancy and without any major underlying pathology, N=5 of which were included for transcriptomics of T cell populations, N=4 for suppression assays, and N=11 for flow cytometry. Baseline characteristics are provided in supplementary table 1. All patients received study information and signed informed consent prior to participation. This study was reviewed and approved by the local Institutional Ethical Review Board of the University Medical Center Utrecht (16-198). One tube of sodium-heparin blood was taken before Caesarean section. After delivery of the neonate and placenta, the placental bed was manually located and two biopsies of the central placental bed from the inner uterine myometrial wall were obtained as previously described.^71^ Additionally, biopsies were taken from the incision site when the placenta was not situated on this part of the uterine wall.

### Lymphocyte isolation

Peripheral blood mononuclear cells (PBMC) were isolated from blood diluted 1:1 with basic medium (RPMI 1640 (Gibco) with Penicillin/Streptomycin (Gibco), L-glutamine (Gibco)), by ficoll-density centrifugation (GE Healthcare-Biosciences, AB). PBMC were washed in basic medium with 2% fetal calf serum (FCS, Biowest) and PBS or staining buffer consisting of cold PBS supplemented with 2% FCS and 0.1% sodium-azide (Severn Biotech Ltd.). The biopsy samples were collected in basic medium supplemented with 10% FCS and minced into pieces of 1 mm^3^ in PBS (Gibco). The biopsies were enzymatically digested with 1 mg/mL collagenase IV (Sigma) in medium for 60 minutes at 37°C in a tube shaker under constant agitation at 120 rpm. To dissolve the remaining biopsy pieces after digestion and remove any remaining lumps, the biopsies were pipetted up and down multiple times and poured over a 100 μm Cell Strainer (BD Falcon). Cells were subsequently washed in staining buffer and filtered through a 70 μm cell strainer and prepared for flow cytometry or flow cytometry-assisted cell sorting.

### Flow cytometry

For flow cytometric experiments without restimulation, PBMC and uterine cells were first incubated in Fixable viability dye eFluor506 (eBioscience) 1:300 in PBS for 20 min at 4°C, and washed in PBS. For surface staining cells were incubated with the antibodies shown in supplementary table 2 for 20 minutes in staining buffer at 4°C, and subsequently washed in the same buffer. Cells were permeabilized with 1 part fixation/permeabilization concentrate and 3 parts fixation/permeabilization diluent (eBioscience) for 30 minutes at 4°C and subsequently incubated overnight with intracellular antibodies (supplementary table 2) in 10x diluted Permeabilization buffer (Perm, eBioscience) 4°C. The next day, cells were washed with Perm and measured on the LSR Fortessa (BD). For intracellular cytokine measurement, PBMC and uterine cells were first incubated with surface staining, washed, and then restimulated with 20 ng/ml phorbol 12-myristate 13-acetate (PMA, Sigma) and 1 μg/ml ionomycin (Sigma) for 4 hours with addition of Monensin (Golgistop, BD Bioscience) during the last 3.5 hours at 37°C. Afterwards, cells were stained with the viability dye, permeabilized, intracellularly stained and measured as described above.

### Flow cytometry-assisted cell sorting

Cells were incubated with surface antibodies (supplementary table 2) for 20 minutes in staining buffer at 4°C, washed in the same buffer and filtered through a 50 μm cell strainer (Filcon, BD). For suppression assays, cells of the CD3^+^CD4^+^CD25^+^CD127^−^ cell population (Tregs) and CD3^+^CD4^+^CD25^−^ cell population (Tconv) were directly sorted into tubes with 500 μL FCS on a FACSAria™ III (BD). For RNA sequencing, 2000 cells of the CD3^+^CD4^+^CD25^+^CD127^−^ cell population (Tregs) and CD3^+^CD4^+^CD25^−^CD45RA^−^ (CD69^+^ from biopsies, CD69^−^ from blood) cell population (Tconv) were sorted into Eppendorfs containing 125 μL PBS. After sorting, 375 μL Trizol LS (Thermo Fisher Scientific) was added to each vial and vials were stored at −80°C until RNA isolation.

### Suppression assays and cytokine measurement

After sorting, peripheral blood and uterine Tregs and Tconv were washed in PBS and resuspended in basic medium with 10% human AB serum (Sanquin). Previously isolated and frozen healthy donor (HC) PBMC were labelled with 2 μM CellTrace Violet (ThermoFisher) as described previously.^72^ Treg or Tconv populations were added to 15.000 HC PBMC at different ratios and cells were co-incubated for 4 days at 37°C. Supernatants were collected for cytokine measurement by multiplex assay before cells were stained with surface antibodies for CD3, CD4 and CD8 as described above and measured on a FACS Canto (BD).

### Whole transcriptome sequencing

For RNA isolation, the vials were thawed at room temperature and 100 μL chloroform was added to each vial. The vials were shaken well and spun down at 12000g for 15 minutes at 4°C. The aqueous phase was transferred into a new tube and RNA was mixed with 1ul of GlycoBlue (Invitrogen) and precipitated with 250 μL isopropanol. Cells were incubated at −20°C for one hour and subsequently spun down at 12000g for 10 minutes. The supernatant was carefully discarded and the RNA pellet was washed twice with 375 μL 75% ethanol. Vials were stored at −80°C until library preparation. Low input RNA sequencing libraries from biological sorted cell population replicates were prepared using the Cel-Seq2 Sample Preparation Protocol^73^ and sequenced as 2 × 75bp paired-end on a NextSeq 500 (Utrecht Sequencing Facility). The reads were demultiplexed and aligned to human cDNA reference using the BWA (0.7.13).^74^ Multiple reads mapping to the same gene with the same unique molecular identifier (UMI, 6bp long) were counted as a single read.

### Data analysis

RNA sequencing data were normalized per million reads per sample. Differentially expressed genes between the cell populations were identified using the DESeq2 package in R 3.5.1 (CRAN), with correction for donor batch (design=~Donor+Cellpop) and input of all genes. Genes with false discovery rate (FDR) adjusted p value (padj)<0.05 and log2(Fold Change)>0.5 or <−0.05 (further annotated as |Log2FC|>0.5) were considered differentially expressed. Principal component analysis (PCA) was performed in DESeq2 based on the constructed model including donor correction. Pathway enrichment analysis was conducted in Toppgene Suite publicly available online portal and pathways with Bonferroni-corrected p-values<0.05 were considered statistically significant.^75^ For heatmap analysis, gene expression was mean-centered and scaled per gene and hierarchical clustering was performed with Ward’s method and Euclidian distance. Gene set enrichment analysis (GSEA^76^) was conducted with Broad Institute software, by 1000 random permutations of the phenotypic subgroups to establish a null distribution of enrichment score, against which a normalized enrichment scores and multiple testing FDR-corrected q values were calculated. Gene sets with an FDR<0.05 were considered significantly enriched. Gene sets were either obtained from provided data in publications or by analyzing raw data using GEO2R (NCBI tool).^77^ An overview of used signatures is provided in supplementary table 3. For flow cytometric data, median fluorescent intensities (MFI) and percentages of positive cells were analyzed in FlowJo (LLC). For graphic representation, data were analyzed in GraphPad Prism (GraphPad Software). To assess significant differences on protein level between groups, Two-way ANOVA with Tukey post hoc test was used and adjusted p values<0.05 were considered statistically significant.

