## Supplementary figure legends for "Human regulatory T cells at the maternal-fetal interface show functional site-specific adaptation with tumor-infiltrating-like features"

**Supplementary figure 1. uTregs derived from the maternal-fetal interface are bona fide maternal Tregs. (A)** Ex vivo frequency of CD25^+^FOXP3^+^ cells among CD4^+^ T cells from blood, maternal-fetal interface (^pb^U) and incision site (^inc^U). **(B)** Sorting strategy for uTregs and uTconv in the uterus. **(C)** Expression of female-specific gene XIST in all sorted T cell subsets. **(D+E)** Suppresssion assay assessing proliferation of healthy CD4^+^ T cells by CTV dilution assay, after 4 days of coculture with healthy donor bTregs, maternal bTregs, or uTregs at a 1:8, 1:4 and 1:2 ratio (n=4).

**Supplementary figure 2. Enrichment of gene signatures of *in vitro* activated Tregs.** Gene set enrichment analysis with genes upregulated in Tregs stimulated *in vitro* with TCR stimulation, IL-2 or TNFα, comparing uTregs and bTregs. ^88–92^ NES = normalized enrichment score.

**Supplementary figure 3. Downregulated pathways shared among uTregs and uTconv.** Pathway analysis (ToppGene Suite) of downregulated pathways shared among uTreg and uTconv. P-values<0.05 after Bonferroni correction were considered significant.

**Supplementary figure 4. uTregs from the maternal-fetal interface share similarities were human Tregs from healthy tissue sites.** Gene set enrichment analysis with published genes which are significantly upregulated in tissue Tregs from healthy skin^15^, colon^14^ or lung^14^ compared to blood Tregs, in uTregs versus bTregs.

**Supplementary figure 5. uTregs from the incision site are bona fide Tregs and have a tissue-resident profile. (A)** GSEA with published Treg signature gene set comparing ^inc^uTreg and ^inc^uTconv.^78^ (**B+C**) Suppresssion assay assessing proliferation of anti-CD3 stimulated (or unstimulated) healthy CD4^+^ T cells by CTV dilution assay **(C)** and cytokine production in the supernatant by multiplex immunoassay **(B)**, after 4 days of coculture with healthy donor bTregs, maternal bTregs, or uTregs at a 1:8, 1:4 and 1:2 ratio. **(D)** Gene expression of CD69. **(E)** Heatmap with published human core tissue-resident genes^4^ in ^inc^uTregs vs bTregs **(F)**. Gene set enrichment analysis (GSEA) with published genes identifying human lung CD4^+^ and CD8^+^ TRM compared to blood memory cells; left panel^93^) and genes upregulated in skin CD4+ TRM compared to blood CD4+ T cells (right panel^15^), in ^inc^uTregs vs bTregs. NES = normalized enrichment score. **(G)** Pathway analysis (ToppGene Suite) with 558 genes upregulated in ^pb^uTregs vs ^inc^uTregs. P-values<0.05 after Bonferroni correction were considered significant. **(H)** GSEA with gene set of upregulated genes in *in vitro* stimulated Tregs.^92^ *NES = normalized enrichment score.*
