## Supplementary figures and images for "Human regulatory T cells at the maternal-fetal interface show functional site-specific adaptation with tumor-infiltrating-like features"

### Supplementary figure 1

#### Supplementary figure 1

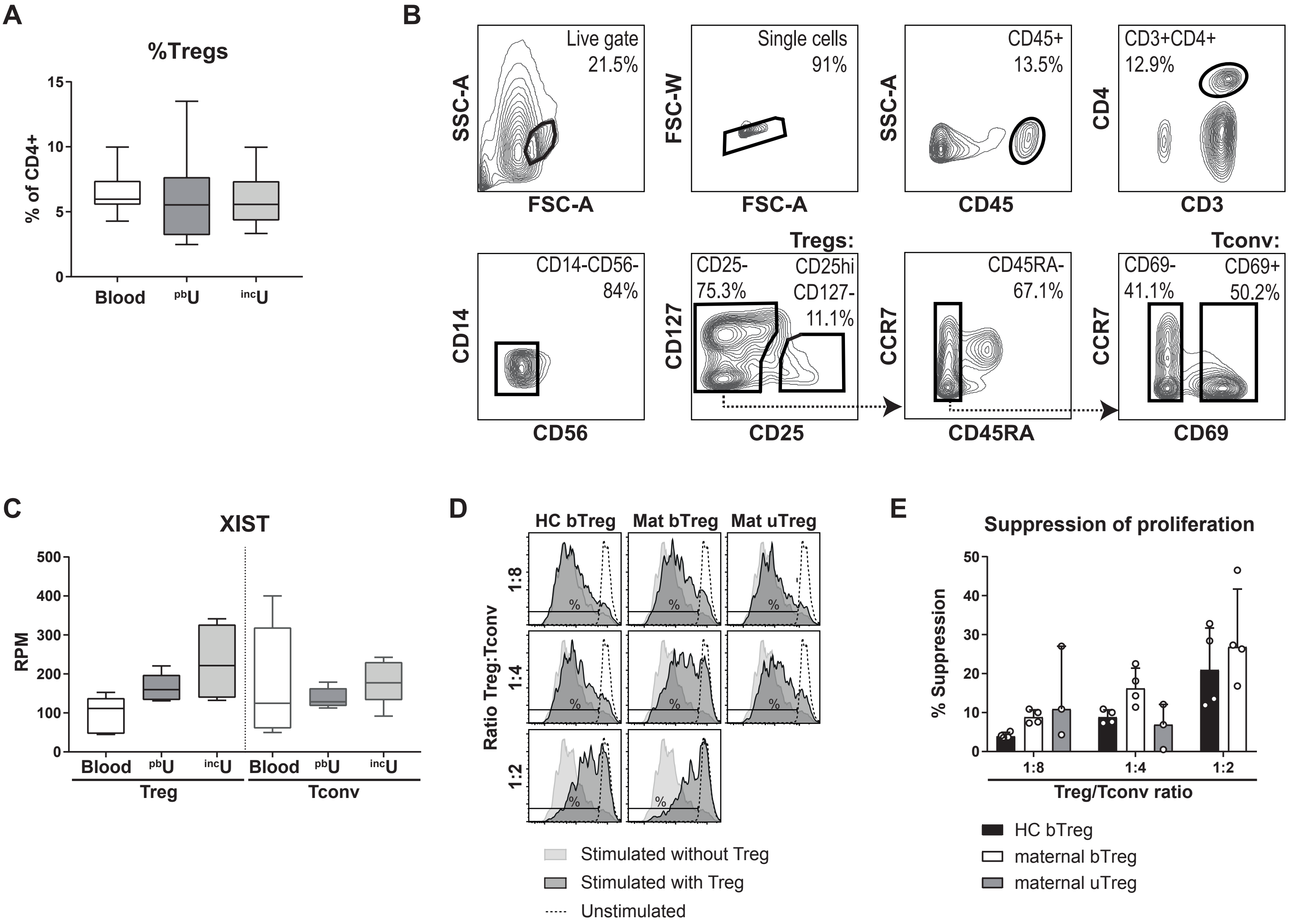

### Supplementary figure 2

##### Supplementary figure 2

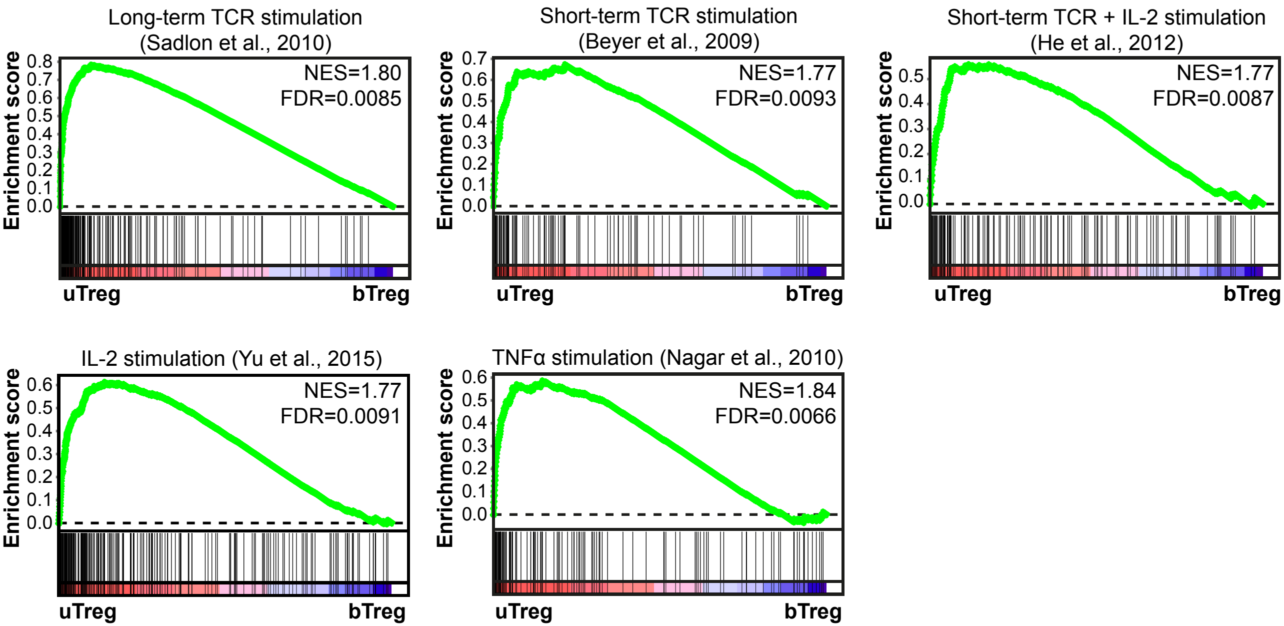

### Supplementary figure 3

##### Supplementary figure 3

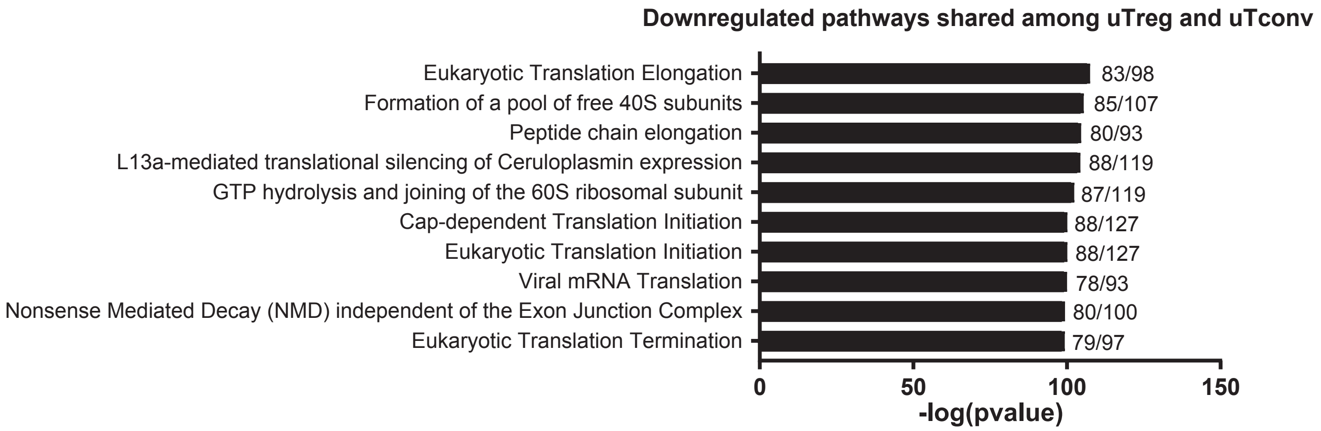

### Supplementary figure 4

##### Supplementary figure 4

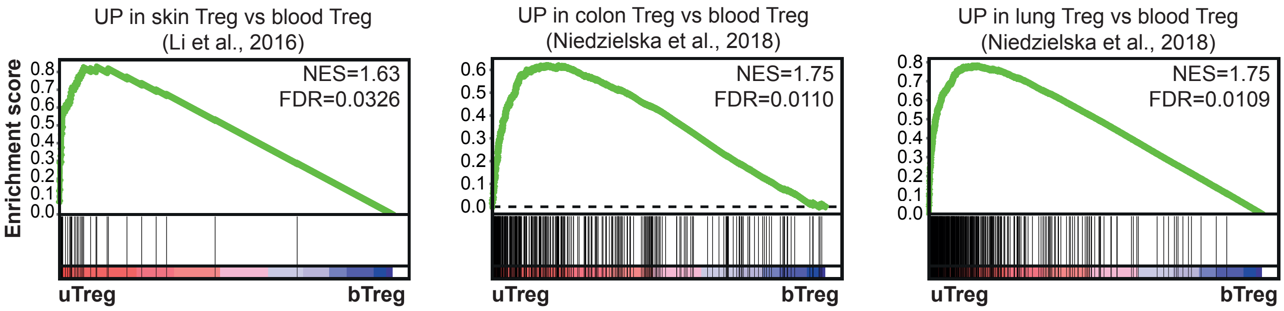

### Supplementary figure 5

#### Supplementary figure 5

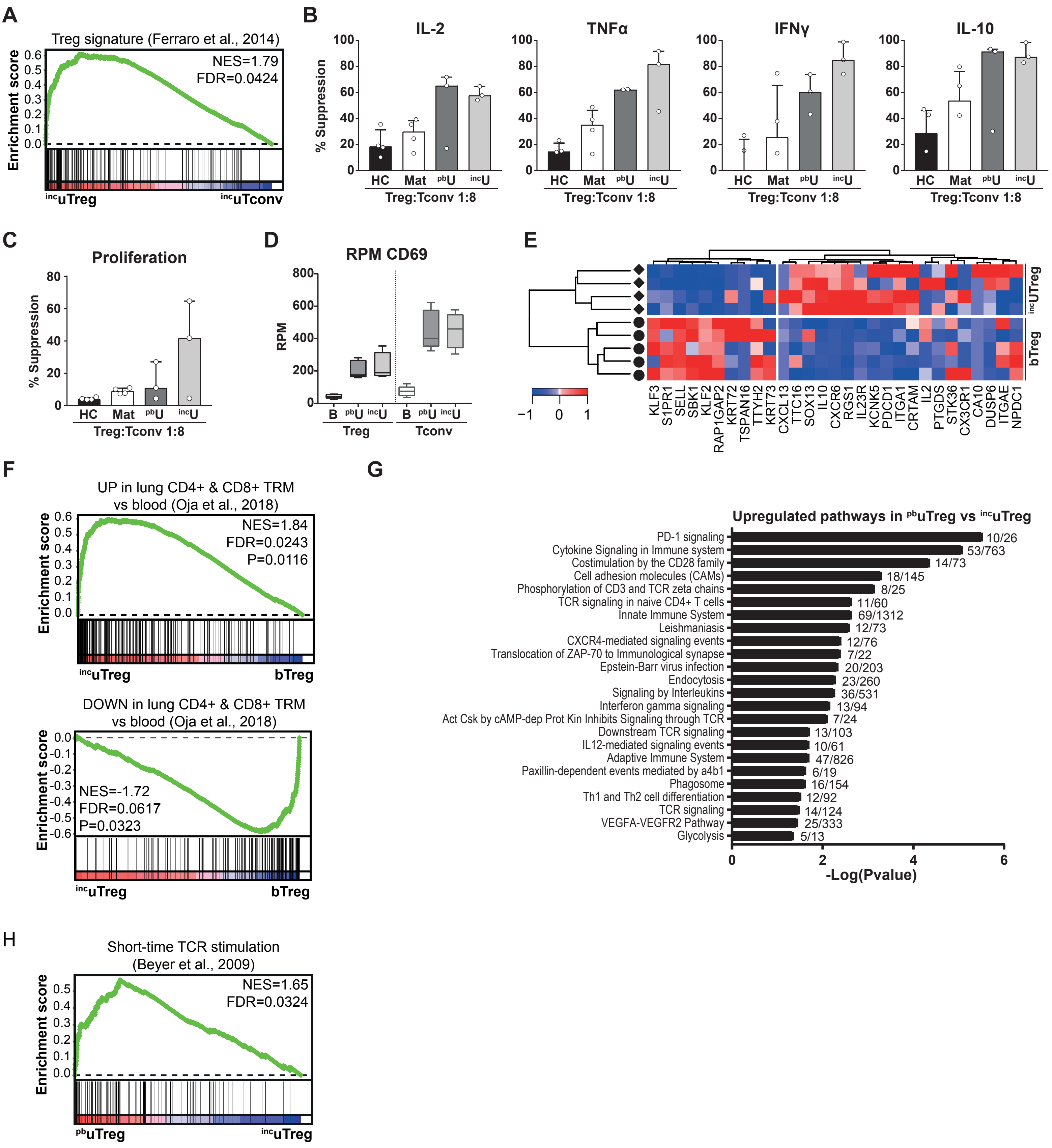
