## Supplementary table 1 for "Human regulatory T cells at the maternal-fetal interface show functional site-specific adaptation with tumor-infiltrating-like features"

**Supplementary table 1. Clinical characteristics of 20 human subjects undergoing caesarian section**

| **Maternal characteristics** |  |
| --- | --- |
| Age (years)*, mean (st.dev)* | 34 (2.3) |
| White ethnicity*, n (%)* | 19 (95%) |
| Gravida (n)*, mean (st.dev)* | 2.3 (1.0) |
| Para (n)*, mean (st.dev)* | 1.0 (0.7) |
| Nulliparous*, n (%)* | 4 (20%) |
| Pregravid BMI (kg/m^2^)*, mean (st.dev)* | 24.4 (5.2) |
| **Neonatal characteristics** |  |
| Gestation age at birth (days), *mean (st.dev)* | 275 (5) |
| Birthweight (grams), *mean (st.dev)* | 3535 (480) |
| Birthweight (percentile), *mean (st.dev)* | 60 (30) |
| FGR (birthweight <p10), *n (%)* | 0 (0%) |
| LGA (birthweight >p95), *n (%)* | 2 (10%) |
| Male sex, *n (%)* | 6 (30%) |
| Apgar at 5 min post partum, *mean (st.dev)* | 8.7 (0.8) |
| Apgar at 10 min post partum, *mean (st.dev)* | 9.6 (0.9) |
