## Supplementary table 2 for "Human regulatory T cells at the maternal-fetal interface show functional site-specific adaptation with tumor-infiltrating-like features"

| **Supplementary table 2. Antibodies used for sorting and flow cytometric analysis** | | | | |  |
| --- | --- | --- | --- | --- | --- |
| **Cell sorting** |  |  |  |  |  |
| *Antibody* | *Fluorochrome* | *Clone* | *Dilution (x)* | *Catalog no* | *Company* |
| CD69 | FITC | FN50 | 25 | 130-113-523 | Miltenyi |
| CCR7 | PE | 3D12 | 25 | 12-1979-42 | eBioscience |
| CD4 | PerCP-Cy5.5 | RPA-T4 | 200 | 2102650 | Sony Biotechnology |
| CD25 | PE-Cy7 | M-A251 | 25 | 557741 | BD |
| CD127 | AF647 | HCD127 | 50 | 2356590 | Sony Biotechnology |
| CD14 | APC-Cy7 | MphiP9 | 200 | 557831 | BD |
| CD56? | PE-CF594 | B159 |  | 562289 | BD |
| CD45RA | PacBlue | HI100 | 200 | 2120590 | Sony Biotechnology |
| CD3 | BV510 | OKT3 | 200 | 317332 | Biolegend |
| CD45RA | BV711 | HI30 | 400 | 304050 | Biolegend |
| **Flow cytometry** |  |  |  |  |  |
| *Surface antibody* | *Fluorochrome* | *Clone* | *Dilution (x)* | *Catalog no* | *Company* |
| CCR5 | FITC | 2D7/CCR5 | 25 | 555992 | BD |
| CCR8 | PE | L263G8 | 100 | 360603 | Biolegend |
| CD134 (OX-40) | PerCP-Cy5.5 | Ber-ACT35 | 50 | 350010 | Biolegend |
| CD137 (4-1BB) | APC | 4B4-1 | 50 | 550890 | BD |
| CD14 | V500 | M5E2 | 100 | 561391 | BD |
| CD25 | BV711 | 2A3 | 50 | 563159 | BD |
| CD3 | APC-eF780 | UCHT1 | 400 | 47-0038-42 | eBioscience |
| CD4 | PE-Cy5 | RPA-T4 | 50 | 555348 | BD |
| CD56 | BV510 | HCD56 | 50 | 2191700 | Sony Biotechnology |
| CD69 | PE-Cy7 | FN50 | 50 | 557745 | BD |
| CXCR3 | BV605 | G025H7 | 12.5 | 2368640 | Sony Biotechnology |
| CXCR5 | PerCP-Cy5.5 | TG2/CXCR5 | 200 | TG2/CXCR5 | AntibodyChain |
| GITR | FITC | #110416 | 25 | FAB689F | R&D |
| HLA-DR | PerCP-Cy5.5 | L243 | 100 | 307630 | Biolegend |
| ICOS | APC | ISA3 | 25 | 17-9948-42 | eBioscience |
| IL18R1 | FITC | H44 | 12.5 | 11-7183-42 | eBioscience |
| PD-1 | APC | MIH4 | 12.5 | 558694 | BD |
| TIGIT | PerCP-eF710 | MBSA43 | 50 | 46-9200-42 | eBioscience |
| *Intracellular antibody* | *Fluorochrome* | *Clone* | *Dilution (x)* | *Catalog no* | *Company* |
| CTLA-4 | PE | BNI3 | 12.5 | 555853 | BD |
| FOXP3 | eFluor450 | PCH101 | 50 | 48-4776-42 | eBioscience |
| Ki67 | AF647 | B56 | 200 | 558615 | BD |
| RORγt | APC | AFKJS-9 | 200 | 17-6988-82 | eBioscience |
| T-bet | PE-CF594 | O4-46 | 25 | 562467 | BD |
| *eBioscience™ Fixable Viability Dye* | *eFluor506* |  | *300* | *65-0866-14* | *ThermoFisher* |
