## Supplementary table 3 for "Human regulatory T cells at the maternal-fetal interface show functional site-specific adaptation with tumor-infiltrating-like features"

| **Supplementary table 3. Gene signatures used for gene set enrichment analysis and overlap with uTreg signature** | | | | | | | | |
| --- | --- | --- | --- | --- | --- | --- | --- | --- |
|  | **Reference** |  | **Geneset origin** | | **Criteria for geneset** | | | |
| *Figure* | *Author, year* | *GEO NCBI* | *Organism* | *Tissue compartment* | *Publ/GEO2R* | *Pvalue/Padj* | *FC* | *Comparison* |
| 1BC, S5A | Ferraro et al., 2014 | - | Human | Peripheral blood | Publication |  |  | Treg signature vs Tconv |
| 1I | Joller et al., 2014 | - | Mouse | Spleen | Publication | P<0.05 | FC>2 | TIGIT+ vs TIGIT- Tregs |
| 2B, S5E | Kumar et al., 2017 | GSE94964 | Human | Lung and spleen | Publication | Padj<0.05 |  | CD69+ vs CD69- (shared among CD4+ and CD8+ from lung and spleen) |
| 2D | Niedzielska et al., 2018 | - | Human | Colon & lung vs peripheral blood | Publication | Padj<0.05 | L2FC>2 | Lung/Colon CD4+ vs blood CD4+ |
| 3C | Li et al., 2016 | GSE74158 | Human | Skin vs peripheral blood | GEO2R | Padj<0.05 |  | Skin CD4+ vs blood CD4+ |
| 3C, S5F | Oja et al., 2018 | - | Human | Lung vs peripheral blood | Publication | Padj<0.05 |  | Shared between lung CD4+ and CD8+ TRM vs Blood TEM |
| 4J | Tan et al., 2016 | - | Mouse | Pancreas prediabetic T1D mice vs spleen | Publication | <0.05 | FC>2 | Splenic CXCR3+ versus CXCR3− Tregs (T-bet+) |
| 5ABC | Dispirito et al., 2018 | - | Mouse | VAT, muscle, Colon vs spleen | Publication | <0.05 | FC>2 | VAT/muscle/colon Tregs vs spleen Tregs |
| 5DEGH | Plitas et al., 2016 | - | Human | Breast cancer vs breast parenchyma vs peripheral blood | Publication | Padj<0.05 |  | UP in breast cancer Treg vs breast cancer Tconv AND UP in breast cancer Treg vs Blood Treg |
| 5DEGH | Toker et al., 2018 | - | Human | Epithelial ovarian cancer (mostly high-grade serous) or melanoma | Publication | Padj<0.1 |  | PD1intICOShi population (Tregs) vs Tconv |
| 5DEGH, 6I | Tirosh et al., 2016 | - | Human | Single cell melanoma | Publication | <0.01 (CD4) <0.05 (CD8) | FC>2 | Meloma Treg vs CD4 Tconv and CD8 Tconv |
| 5DEGH, 6I | Zheng et al., 2017 | - | Human | Hepatocellular carcinoma single cell seq | Publ S3 Cluster | Padj<0.05 | FC>2 | Genes specific for Treg cluster identified by single cell seq |
| 5DEGH, 6I | De Simone et al., 2016 | - | Human | Colon and lung cancer vs colon and lung parenchyma vs blood vs Tconv | Publication |  |  | UP in TITR vs parenchyma vs blood vs Tconv |
| 5DEGH, 6I | Pacella et al., 2018 | - | Human | Liver cirrhosis and tumor (CT) or from the peripheral blood (PB) of patients with chronic HCV infection and hepatocellular carcinoma (HCC) | Publication |  | FC>2 | Upregulated in Treg CT OX40+ versus Treg PB OX40-, and not in Treg CT OX40-, Treg PB OX40+ and all Tconv counteparts |
| 5DEGH, 6I | Magnuson et al., 2018 | - | Human | Colon cancer vs healthy colon | Publication |  |  | Tumor-Treg specific in mouse and human and correlation with FOXP3 expression |
| 5DF | Li et al., 2016 | GSE74158 | Human | Skin vs peripheral blood | GEO2R | Padj<0.05 |  | UP in skin Treg vs skin Tconv AND UP in skin Treg vs Blood Treg |
| 5DF | Niedzielska et al., 2018 | - | Human | Colon & lung vs peripheral blood | Publication | Padj<0.05 | L2FC>2 | UP in lung/colong Treg vs lung/colon Tconv AND UP in lung/colon Treg vs Blood Treg |
| 5IJ, 6J | Plitas et al., 2016 | - | Human | Breast cancer vs breast parenchyma | Publication | Padj<0.05 |  | UP in Breast cancer Tregs vs healthy breast parenchyma Tregs |
| 5IJ, 6J | Magnuson et al., 2018 | - | Human | Colon cancer vs healthy colon | Publication |  |  | Mean overall fold change, or high fold change in at least two patients. Mean FoldChange > 2 and nominal p.value <0.02, or FoldChange >4 (UP), or <0.5 and <0.25 (DN). |
| S2 | Nagar et al., 2010 | GSE18893 | Human | Peripheral blood | GEO2R | P<0.05 | L2FC>0.5 | 50ng/mL TNFa stim vs unstim sorted Tregs (2 and 24 hrs combined in GEO2R) |
| S2 | Yu et al., 2015 | GSE49817 | Human | Peripheral blood | GEO2R | Padj<0.05 | UP FC>1.5, DOWN all FC | Sorted Treg stimulated with IL-2 24 hrsvs unstimulated Tregs |
| S2 | He et al., 2012 | GSE11292 | Human | Peripheral blood | GEO2R | P<0.01 | FC>10 | Sorted Treg stim with anti-CD3/CD28 and IL-2 for 3 hours vs unstimulated Tregs |
| S2 | Sadlon et al., 2010 | GSE20934 | Human | Cord blood | GEO2R | P<0.05 | FC>2 | Cord blood Treg expanded with dynabeads for 8 days + PMA/ionomycin vs resting Tregs at day 4 |
| S2, S5H | Beyer et al., 2009 | GSE16835 | Human | Peripheral blood | GEO2R | Padj<0.05 | FC>3 | Tregs undergoing short time stimulation with anti-CD3/CD28 vs ex vivo Treg |
| S4 | Li et al., 2016 | GSE74158 | Human | Skin vs peripheral blood | GEO2R | Padj<0.05 |  | Skin Treg vs blood Treg |
| S4 | Niedzielska et al., 2018 | - | Human | Colon & lung vs peripheral blood | Publication | Padj<0.05 | L2FC>2 | Lung/Colon Treg vs blood Treg |
