## Supplementary table 4 for "Human regulatory T cells at the maternal-fetal interface show functional site-specific adaptation with tumor-infiltrating-like features"

| **Supplementary table 4. Upregulated and downregulated genes in the uTreg-specific core signature** | | | | | | | | |
| --- | --- | --- | --- | --- | --- | --- | --- | --- |
| **uTreg core UP** | | | **UP vs bTreg** | | | **UP vs uTconv** | | |
| *Gene* | *Ensembl* | *baseMean* | *Log2(FC)* | *Pvalue* | *Padj* | *Log2(FC)* | *Pvalue* | *Padj* |
| **AC006064.4** | ENSG00000269968 | 153.4 | 1.22 | 1.09E-07 | 2.43E-06 | 0.75 | 9.81E-04 | 4.01E-02 |
| **AC017002.1** | ENSG00000240350 | 24.1 | 2.69 | 6.15E-09 | 1.78E-07 | 3.09 | 3.19E-11 | 2.93E-08 |
| **AC080038.1** | ENSG00000011028 | 5.4 | 3.80 | 1.31E-05 | 1.87E-04 | 3.12 | 6.80E-05 | 5.38E-03 |
| **AC126603.1** | ENSG00000258628 | 76.9 | 1.49 | 1.00E-09 | 3.39E-08 | 1.09 | 4.15E-06 | 5.25E-04 |
| **AC132825.2** | ENSG00000243655 | 163.7 | 0.89 | 5.90E-05 | 7.19E-04 | 0.73 | 8.51E-04 | 3.65E-02 |
| **AC147651.4** | ENSG00000237181 | 7.6 | 1.49 | 3.93E-04 | 3.73E-03 | 1.78 | 2.30E-05 | 2.23E-03 |
| **ACP5** | ENSG00000102575 | 28.6 | 1.58 | 2.39E-05 | 3.21E-04 | 1.92 | 2.74E-07 | 5.31E-05 |
| **ADAMTS5** | ENSG00000154736 | 13.3 | 2.94 | 2.54E-07 | 5.30E-06 | 2.21 | 1.78E-05 | 1.85E-03 |
| **ADPRH** | ENSG00000144843 | 8.3 | 1.56 | 7.12E-05 | 8.48E-04 | 1.92 | 1.15E-06 | 1.81E-04 |
| **ADRM1** | ENSG00000130706 | 66.0 | 0.83 | 3.27E-04 | 3.20E-03 | 0.75 | 9.77E-04 | 4.00E-02 |
| **AGTRAP** | ENSG00000177674 | 44.8 | 1.36 | 7.07E-09 | 2.01E-07 | 0.84 | 1.67E-04 | 1.10E-02 |
| **AK4** | ENSG00000162433 | 5.6 | 4.35 | 6.96E-08 | 1.62E-06 | 1.91 | 4.74E-04 | 2.40E-02 |
| **AKIRIN2** | ENSG00000135334 | 44.4 | 1.48 | 8.04E-07 | 1.51E-05 | 1.31 | 8.68E-06 | 1.02E-03 |
| **AL390719.1** | ENSG00000217801 | 15.6 | 1.55 | 2.11E-07 | 4.49E-06 | 1.05 | 7.25E-05 | 5.67E-03 |
| **ARHGAP11B** | ENSG00000187951 | 13.7 | 1.33 | 2.30E-05 | 3.09E-04 | 0.97 | 8.87E-04 | 3.75E-02 |
| **ARID5B** | ENSG00000150347 | 203.1 | 0.88 | 3.06E-06 | 5.11E-05 | 1.04 | 3.32E-08 | 1.06E-05 |
| **ARL3** | ENSG00000138175 | 26.3 | 0.93 | 6.30E-04 | 5.58E-03 | 0.88 | 7.74E-04 | 3.39E-02 |
| **ASMTL** | ENSG00000169093 | 29.1 | 0.73 | 4.78E-03 | 3.08E-02 | 0.84 | 1.10E-03 | 4.28E-02 |
| **ATP1B3** | ENSG00000069849 | 166.3 | 1.01 | 7.06E-08 | 1.64E-06 | 0.73 | 7.58E-05 | 5.89E-03 |
| **ATP6V1B2** | ENSG00000147416 | 79.2 | 0.87 | 3.09E-06 | 5.15E-05 | 0.60 | 9.05E-04 | 3.80E-02 |
| **B3GAT1** | ENSG00000109956 | 8.9 | 3.45 | 1.28E-08 | 3.49E-07 | 3.19 | 2.59E-08 | 8.67E-06 |
| **B4GALT5** | ENSG00000158470 | 43.0 | 1.38 | 4.06E-10 | 1.47E-08 | 1.07 | 3.57E-07 | 6.56E-05 |
| **BABAM2** | ENSG00000158019 | 40.5 | 1.07 | 4.84E-05 | 6.05E-04 | 1.10 | 2.14E-05 | 2.14E-03 |
| **BATF** | ENSG00000156127 | 130.8 | 2.52 | 5.88E-16 | 6.25E-14 | 2.33 | 4.39E-14 | 8.43E-11 |
| **BCL2L11** | ENSG00000153094 | 37.1 | 0.94 | 4.88E-05 | 6.10E-04 | 1.42 | 1.11E-09 | 6.88E-07 |
| **BPGM** | ENSG00000172331 | 18.7 | 1.47 | 6.12E-06 | 9.56E-05 | 1.32 | 2.40E-05 | 2.31E-03 |
| **BST2** | ENSG00000130303 | 59.7 | 1.66 | 7.32E-12 | 3.62E-10 | 0.82 | 3.35E-04 | 1.85E-02 |
| **BTG3** | ENSG00000154640 | 97.0 | 2.33 | 1.13E-21 | 2.79E-19 | 0.91 | 7.65E-05 | 5.92E-03 |
| **C15orf39** | ENSG00000167173 | 14.3 | 1.01 | 2.49E-03 | 1.79E-02 | 1.24 | 1.78E-04 | 1.14E-02 |
| **CAMK1** | ENSG00000134072 | 22.1 | 2.67 | 2.90E-15 | 2.75E-13 | 1.75 | 6.98E-09 | 3.01E-06 |
| **CAVIN3** | ENSG00000170955 | 17.0 | 3.95 | 5.62E-08 | 1.33E-06 | 2.07 | 4.95E-04 | 2.46E-02 |
| **CCDC71L** | ENSG00000253276 | 15.8 | 0.98 | 6.83E-03 | 4.13E-02 | 1.38 | 1.46E-04 | 9.89E-03 |
| **CCL3** | ENSG00000277632 | 4.7 | 3.15 | 5.15E-05 | 6.40E-04 | 2.37 | 6.85E-04 | 3.11E-02 |
| **CCL3** | ENSG00000278567 | 5.1 | 3.74 | 9.33E-06 | 1.39E-04 | 3.49 | 1.01E-05 | 1.16E-03 |
| **CCR1** | ENSG00000163823 | 12.1 | 8.33 | 3.68E-16 | 4.06E-14 | 2.05 | 2.57E-05 | 2.43E-03 |
| **CD74** | ENSG00000019582 | 1077.4 | 1.00 | 2.59E-04 | 2.62E-03 | 1.55 | 1.62E-08 | 5.90E-06 |
| **CD80** | ENSG00000121594 | 4.0 | 3.26 | 2.86E-06 | 4.81E-05 | 3.41 | 5.52E-07 | 9.56E-05 |
| **CDKN1C** | ENSG00000273707 | 13.9 | 2.64 | 5.31E-06 | 8.39E-05 | 1.89 | 3.50E-04 | 1.91E-02 |
| **CDKN2A** | ENSG00000147889 | 52.1 | 2.47 | 8.83E-14 | 6.42E-12 | 1.10 | 5.31E-04 | 2.59E-02 |
| **CEBPB** | ENSG00000172216 | 63.7 | 1.73 | 9.23E-12 | 4.48E-10 | 0.89 | 2.17E-04 | 1.32E-02 |
| **CFAP20** | ENSG00000070761 | 85.8 | 1.21 | 3.41E-07 | 6.90E-06 | 0.87 | 1.62E-04 | 1.07E-02 |
| **CGA** | ENSG00000135346 | 4.2 | 6.00 | 3.41E-07 | 6.90E-06 | 4.78 | 3.00E-06 | 4.19E-04 |
| **CHST11** | ENSG00000171310 | 117.7 | 0.86 | 2.00E-05 | 2.75E-04 | 0.87 | 1.22E-05 | 1.36E-03 |
| **CHSY1** | ENSG00000131873 | 39.9 | 1.12 | 1.22E-07 | 2.71E-06 | 0.75 | 1.95E-04 | 1.22E-02 |
| **COL9A2** | ENSG00000049089 | 10.0 | 3.83 | 3.30E-10 | 1.22E-08 | 2.68 | 7.45E-07 | 1.26E-04 |
| **COMT** | ENSG00000093010 | 62.9 | 0.90 | 9.89E-08 | 2.23E-06 | 0.82 | 5.96E-07 | 1.02E-04 |
| **COX17** | ENSG00000138495 | 95.7 | 0.90 | 2.08E-06 | 3.61E-05 | 0.76 | 4.03E-05 | 3.52E-03 |
| **CRADD** | ENSG00000169372 | 6.6 | 2.29 | 3.04E-05 | 3.96E-04 | 1.95 | 1.28E-04 | 8.97E-03 |
| **CSF1** | ENSG00000184371 | 48.9 | 3.53 | 4.64E-22 | 1.22E-19 | 1.33 | 5.67E-05 | 4.59E-03 |
| **CTLA4** | ENSG00000163599 | 91.2 | 1.89 | 3.58E-09 | 1.09E-07 | 2.84 | 1.39E-18 | 5.85E-15 |
| **CTNNA1** | ENSG00000044115 | 22.5 | 1.39 | 2.76E-07 | 5.70E-06 | 0.83 | 8.39E-04 | 3.62E-02 |
| **CTSC** | ENSG00000109861 | 155.9 | 2.06 | 1.81E-16 | 2.11E-14 | 1.28 | 1.93E-07 | 4.08E-05 |
| **CXCR6** | ENSG00000172215 | 65.7 | 3.97 | 9.30E-28 | 4.08E-25 | 1.34 | 4.86E-05 | 4.11E-03 |
| **CXorf40A** | ENSG00000197620 | 33.9 | 0.96 | 8.49E-06 | 1.27E-04 | 0.98 | 2.66E-06 | 3.75E-04 |
| **CXorf40B** | ENSG00000197021 | 22.0 | 0.98 | 3.29E-04 | 3.22E-03 | 1.26 | 3.42E-06 | 4.55E-04 |
| **DNAJC12** | ENSG00000108176 | 7.3 | 2.82 | 3.85E-05 | 4.92E-04 | 2.13 | 8.35E-04 | 3.61E-02 |
| **DPYSL2** | ENSG00000092964 | 37.6 | 1.04 | 5.08E-04 | 4.65E-03 | 1.42 | 2.10E-06 | 3.07E-04 |
| **DUSP10** | ENSG00000143507 | 32.9 | 2.91 | 2.01E-25 | 7.34E-23 | 1.13 | 8.16E-07 | 1.37E-04 |
| **DYNLL1** | ENSG00000088986 | 347.0 | 0.92 | 4.14E-06 | 6.68E-05 | 0.70 | 4.83E-04 | 2.43E-02 |
| **EIF2AK3** | ENSG00000172071 | 11.5 | 1.65 | 3.04E-06 | 5.08E-05 | 1.01 | 1.02E-03 | 4.06E-02 |
| **ELL2** | ENSG00000118985 | 12.2 | 2.11 | 5.86E-06 | 9.19E-05 | 1.40 | 1.17E-03 | 4.55E-02 |
| **ENO1** | ENSG00000074800 | 971.5 | 1.54 | 1.27E-11 | 5.98E-10 | 0.74 | 1.08E-03 | 4.24E-02 |
| **ENTPD1** | ENSG00000138185 | 72.2 | 0.50 | 2.68E-03 | 1.91E-02 | 0.88 | 1.39E-07 | 3.13E-05 |
| **ERI1** | ENSG00000104626 | 26.5 | 1.43 | 6.70E-09 | 1.91E-07 | 1.03 | 9.45E-06 | 1.10E-03 |
| **ETS2** | ENSG00000157557 | 17.9 | 2.36 | 1.88E-07 | 4.04E-06 | 2.21 | 4.31E-07 | 7.77E-05 |
| **ETV7** | ENSG00000010030 | 27.6 | 1.40 | 4.05E-05 | 5.16E-04 | 1.56 | 3.97E-06 | 5.05E-04 |
| **EVA1B** | ENSG00000142694 | 12.9 | 1.21 | 1.54E-03 | 1.19E-02 | 1.62 | 2.69E-05 | 2.53E-03 |
| **FABP5** | ENSG00000164687 | 21.0 | 1.72 | 2.55E-06 | 4.35E-05 | 1.66 | 3.21E-06 | 4.34E-04 |
| **FAM3C** | ENSG00000196937 | 13.2 | 2.39 | 7.45E-11 | 3.09E-09 | 1.56 | 1.21E-06 | 1.88E-04 |
| **FGL2** | ENSG00000127951 | 13.2 | 1.50 | 7.55E-06 | 1.14E-04 | 1.24 | 8.51E-05 | 6.52E-03 |
| **FKBP1A** | ENSG00000088832 | 236.1 | 0.94 | 1.02E-05 | 1.49E-04 | 0.81 | 1.26E-04 | 8.83E-03 |
| **FKBP1C** | ENSG00000198225 | 18.5 | 0.95 | 3.00E-04 | 2.98E-03 | 1.29 | 9.63E-07 | 1.58E-04 |
| **FOXB1** | ENSG00000171956 | 3.2 | 2.44 | 8.75E-04 | 7.40E-03 | 3.48 | 1.26E-05 | 1.40E-03 |
| **FUCA2** | ENSG00000001036 | 19.3 | 1.52 | 2.65E-07 | 5.51E-06 | 0.97 | 3.56E-04 | 1.92E-02 |
| **GADD45A** | ENSG00000116717 | 36.1 | 2.38 | 1.94E-09 | 6.19E-08 | 1.84 | 1.71E-06 | 2.60E-04 |
| **GADD45G** | ENSG00000130222 | 17.4 | 6.43 | 1.48E-15 | 1.47E-13 | 1.88 | 3.63E-06 | 4.70E-04 |
| **GAPDH** | ENSG00000111640 | 955.4 | 1.30 | 1.20E-09 | 4.00E-08 | 0.76 | 3.52E-04 | 1.91E-02 |
| **GEM** | ENSG00000164949 | 10.8 | 8.39 | 1.33E-11 | 6.23E-10 | 5.29 | 3.64E-08 | 1.15E-05 |
| **GLUD1** | ENSG00000148672 | 91.8 | 0.69 | 2.18E-04 | 2.26E-03 | 0.68 | 1.88E-04 | 1.19E-02 |
| **GPAT3** | ENSG00000138678 | 6.8 | 2.33 | 3.17E-06 | 5.26E-05 | 1.48 | 4.98E-04 | 2.47E-02 |
| **GPR137B** | ENSG00000077585 | 15.1 | 2.38 | 7.35E-13 | 4.49E-11 | 0.83 | 1.07E-03 | 4.21E-02 |
| **GSTO1** | ENSG00000148834 | 85.1 | 0.80 | 2.46E-04 | 2.50E-03 | 0.74 | 6.33E-04 | 2.97E-02 |
| **HAVCR2** | ENSG00000135077 | 25.3 | 1.40 | 1.94E-07 | 4.17E-06 | 0.85 | 6.14E-04 | 2.89E-02 |
| **HES1** | ENSG00000114315 | 12.2 | 5.22 | 7.09E-08 | 1.65E-06 | 3.65 | 3.14E-07 | 5.87E-05 |
| **HLA-A** | ENSG00000223980 | 729.5 | 0.70 | 1.28E-03 | 1.02E-02 | 0.75 | 5.52E-04 | 2.66E-02 |
| **HLA-A** | ENSG00000224320 | 848.3 | 0.64 | 1.12E-03 | 9.07E-03 | 0.67 | 6.20E-04 | 2.91E-02 |
| **HLA-A** | ENSG00000227715 | 787.4 | 0.66 | 1.87E-03 | 1.41E-02 | 0.71 | 8.26E-04 | 3.58E-02 |
| **HLA-A** | ENSG00000235657 | 781.5 | 0.63 | 3.15E-03 | 2.18E-02 | 0.69 | 1.15E-03 | 4.48E-02 |
| **HLA-DQB1** | ENSG00000225824 | 33.0 | 1.33 | 9.31E-04 | 7.78E-03 | 1.32 | 8.50E-04 | 3.65E-02 |
| **HLA-DRA** | ENSG00000204287 | 12.0 | 1.31 | 7.35E-03 | 4.37E-02 | 2.75 | 1.18E-07 | 2.78E-05 |
| **HLA-DRA** | ENSG00000227993 | 11.6 | 1.41 | 5.81E-03 | 3.61E-02 | 3.35 | 2.57E-09 | 1.39E-06 |
| **HLA-DRA** | ENSG00000228987 | 13.6 | 1.56 | 1.52E-03 | 1.18E-02 | 3.34 | 4.79E-10 | 3.27E-07 |
| **HLA-DRA** | ENSG00000234794 | 13.2 | 1.88 | 3.23E-05 | 4.18E-04 | 3.94 | 6.25E-14 | 1.10E-10 |
| **HLA-DRB1** | ENSG00000228080 | 111.4 | 1.54 | 3.13E-03 | 2.17E-02 | 2.75 | 2.70E-07 | 5.27E-05 |
| **HLA-DRB1** | ENSG00000229074 | 113.7 | 1.84 | 3.53E-04 | 3.42E-03 | 2.88 | 7.09E-08 | 1.92E-05 |
| **HLA-DRB4** | ENSG00000227357 | 20.9 | 1.55 | 1.97E-03 | 1.48E-02 | 2.20 | 1.43E-05 | 1.55E-03 |
| **HLA-DRB4** | ENSG00000231021 | 21.3 | 1.88 | 7.43E-05 | 8.79E-04 | 2.57 | 8.73E-08 | 2.25E-05 |
| **HNRNPLL** | ENSG00000143889 | 87.6 | 0.79 | 1.13E-04 | 1.26E-03 | 0.71 | 4.60E-04 | 2.35E-02 |
| **IFT27** | ENSG00000100360 | 21.3 | 0.70 | 5.11E-03 | 3.25E-02 | 0.90 | 2.45E-04 | 1.45E-02 |
| **IGFLR1** | ENSG00000126246 | 59.4 | 0.81 | 1.88E-05 | 2.61E-04 | 1.10 | 4.67E-09 | 2.19E-06 |
| **IKZF4** | ENSG00000123411 | 26.2 | 0.81 | 6.43E-03 | 3.93E-02 | 1.53 | 5.39E-07 | 9.42E-05 |
| **IL10** | ENSG00000136634 | 28.3 | 3.05 | 8.34E-10 | 2.84E-08 | 2.56 | 1.05E-07 | 2.55E-05 |
| **IL1R1** | ENSG00000115594 | 22.0 | 1.47 | 9.82E-06 | 1.45E-04 | 1.19 | 1.91E-04 | 1.20E-02 |
| **IL1R2** | ENSG00000115590 | 12.2 | 4.22 | 7.54E-04 | 6.52E-03 | 4.25 | 6.07E-04 | 2.88E-02 |
| **IL1RAP** | ENSG00000196083 | 18.1 | 0.87 | 4.66E-03 | 3.01E-02 | 0.97 | 1.31E-03 | 4.91E-02 |
| **IL1RN** | ENSG00000136689 | 5.1 | 3.51 | 6.95E-05 | 8.31E-04 | 4.56 | 1.72E-06 | 2.60E-04 |
| **IL2RA** | ENSG00000134460 | 114.7 | 0.98 | 4.67E-05 | 5.87E-04 | 3.11 | 2.43E-35 | 5.14E-31 |
| **IL2RB** | ENSG00000100385 | 224.5 | 1.96 | 7.34E-12 | 3.62E-10 | 1.22 | 1.76E-05 | 1.84E-03 |
| **ITGAM** | ENSG00000169896 | 19.0 | 3.14 | 5.08E-08 | 1.22E-06 | 3.07 | 5.50E-08 | 1.59E-05 |
| **JMJD4** | ENSG00000081692 | 43.8 | 1.02 | 1.03E-06 | 1.88E-05 | 0.66 | 8.98E-04 | 3.79E-02 |
| **JOSD2** | ENSG00000161677 | 29.9 | 1.14 | 9.96E-06 | 1.47E-04 | 0.85 | 5.78E-04 | 2.76E-02 |
| **KAT2B** | ENSG00000114166 | 31.9 | 1.39 | 4.80E-08 | 1.16E-06 | 0.81 | 6.97E-04 | 3.12E-02 |
| **KCNK5** | ENSG00000164626 | 5.1 | 4.87 | 3.11E-09 | 9.64E-08 | 2.59 | 4.64E-07 | 8.23E-05 |
| **KDM2A** | ENSG00000173120 | 123.4 | 1.05 | 7.55E-07 | 1.42E-05 | 0.69 | 9.00E-04 | 3.79E-02 |
| **KLHL2** | ENSG00000109466 | 12.8 | 1.18 | 4.52E-04 | 4.22E-03 | 1.12 | 5.40E-04 | 2.61E-02 |
| **LAG3** | ENSG00000089692 | 25.1 | 4.17 | 4.64E-17 | 6.07E-15 | 1.59 | 1.61E-04 | 1.07E-02 |
| **LAPTM4B** | ENSG00000104341 | 47.7 | 1.76 | 1.25E-12 | 7.21E-11 | 0.80 | 5.00E-04 | 2.47E-02 |
| **LAYN** | ENSG00000204381 | 10.0 | 2.84 | 1.24E-08 | 3.38E-07 | 3.48 | 1.82E-11 | 1.83E-08 |
| **LGALS1** | ENSG00000100097 | 340.4 | 2.90 | 6.11E-17 | 7.82E-15 | 1.39 | 5.08E-05 | 4.21E-03 |
| **LGALS3** | ENSG00000131981 | 197.5 | 2.00 | 7.97E-14 | 5.84E-12 | 1.26 | 1.72E-06 | 2.60E-04 |
| **LINC01943** | ENSG00000280721 | 29.5 | 2.91 | 2.85E-10 | 1.08E-08 | 2.56 | 1.46E-08 | 5.56E-06 |
| **LINC02195** | ENSG00000236481 | 6.4 | 2.65 | 4.25E-07 | 8.39E-06 | 4.54 | 3.45E-12 | 4.29E-09 |
| **LRRC32** | ENSG00000137507 | 8.4 | 2.49 | 3.39E-04 | 3.31E-03 | 4.27 | 3.07E-08 | 1.01E-05 |
| **LTA** | ENSG00000231408 | 9.2 | 2.43 | 9.72E-08 | 2.20E-06 | 1.33 | 3.39E-04 | 1.87E-02 |
| **MAF** | ENSG00000178573 | 290.5 | 0.98 | 1.98E-06 | 3.45E-05 | 1.07 | 2.51E-07 | 4.95E-05 |
| **MAPKAPK3** | ENSG00000114738 | 62.8 | 1.97 | 5.20E-16 | 5.60E-14 | 0.72 | 1.28E-03 | 4.86E-02 |
| **MAST4** | ENSG00000069020 | 181.9 | 1.25 | 2.74E-11 | 1.23E-09 | 0.96 | 1.79E-07 | 3.88E-05 |
| **MB21D2** | ENSG00000180611 | 19.9 | 3.33 | 6.67E-11 | 2.78E-09 | 1.68 | 1.70E-04 | 1.11E-02 |
| **MCM6** | ENSG00000076003 | 29.5 | 0.77 | 7.50E-04 | 6.49E-03 | 0.95 | 2.43E-05 | 2.32E-03 |
| **MGST2** | ENSG00000085871 | 15.3 | 0.87 | 1.58E-03 | 1.22E-02 | 1.65 | 1.19E-08 | 4.67E-06 |
| **MIF-AS1** | ENSG00000218537 | 60.7 | 0.79 | 2.99E-04 | 2.97E-03 | 0.73 | 6.10E-04 | 2.88E-02 |
| **MTREX** | ENSG00000039123 | 91.6 | 2.46 | 1.62E-14 | 1.38E-12 | 1.68 | 7.36E-08 | 1.94E-05 |
| **MT-RNR1** | ENSG00000211459 | 4450.1 | 1.73 | 1.53E-20 | 3.19E-18 | 0.60 | 1.28E-03 | 4.86E-02 |
| **MTRNR2L1** | ENSG00000256618 | 2315.5 | 1.43 | 4.17E-16 | 4.53E-14 | 0.60 | 6.77E-04 | 3.09E-02 |
| **MTRNR2L10** | ENSG00000256045 | 141.2 | 1.65 | 1.64E-13 | 1.14E-11 | 0.69 | 1.24E-03 | 4.76E-02 |
| **MYO1E** | ENSG00000157483 | 8.8 | 3.20 | 1.51E-11 | 6.97E-10 | 2.40 | 1.67E-09 | 9.51E-07 |
| **MYO7A** | ENSG00000137474 | 10.7 | 1.67 | 1.49E-04 | 1.62E-03 | 1.90 | 1.54E-05 | 1.64E-03 |
| **NAB1** | ENSG00000138386 | 13.0 | 2.29 | 1.79E-09 | 5.76E-08 | 1.86 | 6.71E-08 | 1.84E-05 |
| **NAMPT** | ENSG00000105835 | 64.2 | 3.21 | 2.78E-59 | 5.23E-56 | 1.81 | 3.15E-29 | 2.22E-25 |
| **NAMPTP1** | ENSG00000229644 | 17.8 | 4.02 | 1.48E-23 | 4.55E-21 | 2.75 | 5.12E-18 | 1.80E-14 |
| **NCF4** | ENSG00000100365 | 43.6 | 0.78 | 1.81E-03 | 1.38E-02 | 1.15 | 4.20E-06 | 5.29E-04 |
| **NDFIP2** | ENSG00000102471 | 26.4 | 3.80 | 5.28E-20 | 1.00E-17 | 1.32 | 9.62E-05 | 7.21E-03 |
| **NDUFV2** | ENSG00000178127 | 124.3 | 0.60 | 1.60E-03 | 1.24E-02 | 0.60 | 1.26E-03 | 4.82E-02 |
| **NFIL3** | ENSG00000165030 | 18.9 | 5.24 | 6.56E-17 | 8.27E-15 | 2.36 | 1.15E-07 | 2.76E-05 |
| **NINJ1** | ENSG00000131669 | 109.4 | 2.23 | 7.31E-17 | 9.16E-15 | 1.18 | 5.11E-06 | 6.31E-04 |
| **NMB** | ENSG00000197696 | 13.6 | 2.25 | 2.23E-09 | 7.02E-08 | 1.61 | 2.46E-06 | 3.51E-04 |
| **NR4A3** | ENSG00000119508 | 39.3 | 7.22 | 7.00E-18 | 9.99E-16 | 0.80 | 5.07E-04 | 2.50E-02 |
| **OGG1** | ENSG00000114026 | 39.8 | 1.04 | 4.55E-07 | 8.94E-06 | 0.66 | 6.41E-04 | 2.98E-02 |
| **OTUD1** | ENSG00000165312 | 33.1 | 1.41 | 5.30E-05 | 6.56E-04 | 1.63 | 2.61E-06 | 3.70E-04 |
| **PARPBP** | ENSG00000185480 | 13.3 | 1.55 | 2.16E-03 | 1.59E-02 | 1.70 | 6.88E-04 | 3.11E-02 |
| **PDCD1** | ENSG00000276977 | 48.4 | 4.09 | 8.79E-26 | 3.41E-23 | 1.16 | 3.57E-04 | 1.92E-02 |
| **PDGFA** | ENSG00000197461 | 11.0 | 1.65 | 1.48E-05 | 2.09E-04 | 1.39 | 9.53E-05 | 7.17E-03 |
| **PELI1** | ENSG00000197329 | 38.8 | 1.31 | 3.30E-06 | 5.42E-05 | 1.96 | 7.41E-12 | 8.24E-09 |
| **PGAM1P8** | ENSG00000255200 | 37.6 | 1.24 | 1.04E-07 | 2.34E-06 | 0.70 | 1.33E-03 | 4.94E-02 |
| **PGK1** | ENSG00000102144 | 412.2 | 1.33 | 3.42E-10 | 1.26E-08 | 0.81 | 1.09E-04 | 7.95E-03 |
| **PGM2L1** | ENSG00000165434 | 96.0 | 1.13 | 1.90E-08 | 5.04E-07 | 0.82 | 2.90E-05 | 2.69E-03 |
| **PHLDA1** | ENSG00000139289 | 302.4 | 4.72 | 2.67E-72 | 1.82E-68 | 0.77 | 1.22E-03 | 4.68E-02 |
| **PHLDA2** | ENSG00000181649 | 10.3 | 4.91 | 2.64E-12 | 1.41E-10 | 1.68 | 6.70E-05 | 5.34E-03 |
| **PHLDA2** | ENSG00000274538 | 10.6 | 4.59 | 1.19E-13 | 8.49E-12 | 1.90 | 2.36E-06 | 3.39E-04 |
| **PHTF2** | ENSG00000006576 | 43.3 | 1.65 | 1.03E-15 | 1.05E-13 | 0.67 | 2.62E-04 | 1.52E-02 |
| **PIGT** | ENSG00000124155 | 43.2 | 0.70 | 1.85E-03 | 1.40E-02 | 0.83 | 1.71E-04 | 1.11E-02 |
| **PIM3** | ENSG00000198355 | 173.1 | 2.55 | 3.74E-17 | 4.95E-15 | 1.04 | 4.16E-04 | 2.19E-02 |
| **PKM** | ENSG00000067225 | 645.7 | 1.18 | 6.06E-06 | 9.47E-05 | 1.22 | 3.07E-06 | 4.27E-04 |
| **PLPP1** | ENSG00000067113 | 55.9 | 3.51 | 7.01E-12 | 3.50E-10 | 2.17 | 1.10E-05 | 1.24E-03 |
| **PMAIP1** | ENSG00000141682 | 41.3 | 2.33 | 4.27E-14 | 3.29E-12 | 1.17 | 4.97E-05 | 4.14E-03 |
| **PMVK** | ENSG00000163344 | 40.2 | 1.04 | 2.94E-06 | 4.94E-05 | 0.69 | 1.26E-03 | 4.80E-02 |
| **PRDM1** | ENSG00000057657 | 221.3 | 0.75 | 7.56E-05 | 8.92E-04 | 1.12 | 3.43E-09 | 1.72E-06 |
| **PRDX1** | ENSG00000117450 | 77.6 | 0.57 | 3.67E-03 | 2.47E-02 | 0.72 | 2.27E-04 | 1.35E-02 |
| **PRNP** | ENSG00000171867 | 62.5 | 1.90 | 2.57E-10 | 9.77E-09 | 0.92 | 1.35E-03 | 4.99E-02 |
| **PSMD1** | ENSG00000173692 | 78.1 | 0.56 | 6.80E-04 | 5.95E-03 | 0.53 | 9.62E-04 | 3.97E-02 |
| **PSMD8** | ENSG00000099341 | 126.4 | 0.51 | 3.59E-04 | 3.47E-03 | 0.59 | 3.06E-05 | 2.83E-03 |
| **PTP4A3** | ENSG00000184489 | 9.6 | 1.72 | 6.52E-04 | 5.75E-03 | 1.80 | 2.79E-04 | 1.59E-02 |
| **PTP4A3** | ENSG00000275575 | 7.4 | 1.66 | 3.52E-03 | 2.40E-02 | 1.98 | 5.20E-04 | 2.55E-02 |
| **PTTG1** | ENSG00000164611 | 34.7 | 1.40 | 1.14E-05 | 1.64E-04 | 1.55 | 1.08E-06 | 1.71E-04 |
| **PXK** | ENSG00000168297 | 14.5 | 1.14 | 1.04E-04 | 1.18E-03 | 0.95 | 6.53E-04 | 3.02E-02 |
| **RAB10** | ENSG00000084733 | 52.4 | 0.92 | 4.96E-07 | 9.67E-06 | 0.60 | 5.66E-04 | 2.70E-02 |
| **RAC1** | ENSG00000136238 | 238.4 | 0.62 | 3.09E-04 | 3.06E-03 | 0.66 | 9.74E-05 | 7.27E-03 |
| **RBKS** | ENSG00000171174 | 7.1 | 2.49 | 5.12E-06 | 8.13E-05 | 1.79 | 2.02E-04 | 1.24E-02 |
| **RCAN2** | ENSG00000172348 | 10.1 | 7.23 | 3.94E-12 | 2.04E-10 | 2.78 | 3.20E-05 | 2.93E-03 |
| **RDH10** | ENSG00000121039 | 21.8 | 2.01 | 4.25E-08 | 1.04E-06 | 2.20 | 1.49E-09 | 8.77E-07 |
| **RHBDD2** | ENSG00000005486 | 119.4 | 1.02 | 3.31E-05 | 4.29E-04 | 0.78 | 1.35E-03 | 4.99E-02 |
| **RHPN2** | ENSG00000131941 | 87.0 | 1.20 | 9.09E-11 | 3.73E-09 | 1.07 | 4.40E-09 | 2.15E-06 |
| **RNF187** | ENSG00000168159 | 79.1 | 0.81 | 2.16E-04 | 2.24E-03 | 0.75 | 5.21E-04 | 2.55E-02 |
| **SAT1** | ENSG00000130066 | 194.6 | 0.79 | 1.03E-03 | 8.44E-03 | 0.94 | 1.07E-04 | 7.82E-03 |
| **SDC4** | ENSG00000124145 | 39.0 | 3.53 | 1.11E-22 | 3.14E-20 | 1.83 | 1.47E-08 | 5.56E-06 |
| **SDF4** | ENSG00000078808 | 115.6 | 0.85 | 9.96E-05 | 1.14E-03 | 0.77 | 3.34E-04 | 1.85E-02 |
| **SEC14L1** | ENSG00000129657 | 64.4 | 0.88 | 1.17E-04 | 1.30E-03 | 0.76 | 7.62E-04 | 3.35E-02 |
| **SETBP1** | ENSG00000152217 | 4.7 | 3.11 | 6.49E-06 | 1.01E-04 | 1.83 | 7.08E-04 | 3.16E-02 |
| **SGMS1** | ENSG00000198964 | 21.7 | 0.86 | 1.02E-03 | 8.41E-03 | 1.19 | 5.65E-06 | 6.90E-04 |
| **SIGLEC17P** | ENSG00000171101 | 15.4 | 5.68 | 5.02E-15 | 4.70E-13 | 3.41 | 3.33E-10 | 2.35E-07 |
| **SIPA1L1** | ENSG00000197555 | 36.4 | 1.50 | 4.21E-09 | 1.25E-07 | 1.07 | 8.86E-06 | 1.03E-03 |
| **SLC16A1** | ENSG00000155380 | 9.0 | 1.59 | 3.33E-05 | 4.31E-04 | 1.49 | 4.48E-05 | 3.86E-03 |
| **SLC27A2** | ENSG00000140284 | 4.6 | 4.87 | 5.27E-07 | 1.02E-05 | 3.45 | 8.25E-06 | 9.79E-04 |
| **SLC5A3** | ENSG00000198743 | 33.6 | 1.96 | 2.69E-16 | 3.06E-14 | 0.69 | 6.94E-04 | 3.12E-02 |
| **SLC7A5** | ENSG00000103257 | 72.9 | 3.63 | 2.77E-23 | 8.34E-21 | 1.45 | 1.83E-05 | 1.88E-03 |
| **SLCO4A1** | ENSG00000101187 | 4.8 | 3.53 | 5.97E-07 | 1.15E-05 | 2.08 | 2.27E-04 | 1.35E-02 |
| **SMOX** | ENSG00000088826 | 6.0 | 4.83 | 5.65E-08 | 1.34E-06 | 2.20 | 1.63E-04 | 1.07E-02 |
| **SNAP47** | ENSG00000143740 | 41.5 | 2.09 | 3.46E-12 | 1.81E-10 | 1.16 | 4.04E-05 | 3.52E-03 |
| **SNX5** | ENSG00000089006 | 76.3 | 0.52 | 2.92E-04 | 2.91E-03 | 0.56 | 5.66E-05 | 4.59E-03 |
| **SNX9** | ENSG00000130340 | 38.7 | 1.64 | 7.13E-07 | 1.35E-05 | 1.38 | 2.01E-05 | 2.04E-03 |
| **SOX4** | ENSG00000124766 | 12.6 | 1.89 | 2.02E-04 | 2.12E-03 | 2.21 | 1.46E-05 | 1.58E-03 |
| **SPATS2L** | ENSG00000196141 | 90.6 | 1.55 | 1.12E-19 | 2.01E-17 | 0.81 | 4.40E-07 | 7.87E-05 |
| **SRGN** | ENSG00000122862 | 1731.7 | 2.71 | 1.15E-46 | 1.36E-43 | 0.69 | 2.21E-04 | 1.33E-02 |
| **SURF4** | ENSG00000148248 | 73.2 | 1.49 | 3.46E-10 | 1.27E-08 | 0.74 | 1.20E-03 | 4.64E-02 |
| **SUSD6** | ENSG00000100647 | 70.5 | 1.14 | 3.47E-06 | 5.68E-05 | 0.80 | 9.72E-04 | 3.99E-02 |
| **SYT11** | ENSG00000132718 | 44.7 | 0.69 | 4.55E-04 | 4.24E-03 | 1.08 | 4.90E-08 | 1.48E-05 |
| **TFRC** | ENSG00000072274 | 78.9 | 1.02 | 3.36E-08 | 8.38E-07 | 1.28 | 3.30E-12 | 4.29E-09 |
| **TGIF2-RAB5IF** | ENSG00000259399 | 8.7 | 0.98 | 7.50E-03 | 4.44E-02 | 1.24 | 6.52E-04 | 3.02E-02 |
| **TMED3** | ENSG00000166557 | 67.3 | 0.59 | 2.95E-04 | 2.94E-03 | 0.77 | 1.49E-06 | 2.30E-04 |
| **TMEM173** | ENSG00000184584 | 225.7 | 0.87 | 6.27E-06 | 9.77E-05 | 0.65 | 6.75E-04 | 3.09E-02 |
| **TNFRSF13B** | ENSG00000240505 | 4.5 | 3.63 | 6.63E-06 | 1.02E-04 | 6.76 | 1.88E-09 | 1.05E-06 |
| **TNFRSF18** | ENSG00000186891 | 58.6 | 4.15 | 3.77E-22 | 1.03E-19 | 2.19 | 3.20E-08 | 1.04E-05 |
| **TNFRSF1B** | ENSG00000028137 | 539.9 | 1.11 | 9.86E-06 | 1.46E-04 | 1.20 | 1.91E-06 | 2.85E-04 |
| **TNFRSF4** | ENSG00000186827 | 95.2 | 3.07 | 7.53E-17 | 9.38E-15 | 2.82 | 1.04E-14 | 2.45E-11 |
| **TNFRSF8** | ENSG00000120949 | 2.8 | 2.44 | 3.66E-03 | 2.47E-02 | 4.56 | 2.11E-05 | 2.12E-03 |
| **TNFRSF9** | ENSG00000049249 | 74.3 | 1.55 | 9.01E-11 | 3.70E-09 | 1.13 | 1.06E-06 | 1.69E-04 |
| **TNIP2** | ENSG00000168884 | 77.4 | 0.95 | 4.76E-04 | 4.41E-03 | 0.92 | 6.80E-04 | 3.09E-02 |
| **TNS3** | ENSG00000136205 | 19.4 | 2.25 | 2.74E-12 | 1.45E-10 | 0.92 | 4.31E-04 | 2.26E-02 |
| **TOX2** | ENSG00000124191 | 14.5 | 2.31 | 1.08E-04 | 1.22E-03 | 2.66 | 8.41E-06 | 9.93E-04 |
| **TP53INP2** | ENSG00000078804 | 15.0 | 4.26 | 1.64E-14 | 1.39E-12 | 1.03 | 9.85E-04 | 4.01E-02 |
| **TPI1** | ENSG00000111669 | 218.1 | 1.50 | 2.84E-08 | 7.24E-07 | 0.99 | 2.46E-04 | 1.45E-02 |
| **TPP1** | ENSG00000166340 | 187.5 | 0.66 | 1.12E-03 | 9.09E-03 | 0.68 | 6.95E-04 | 3.12E-02 |
| **TRAF1** | ENSG00000056558 | 64.5 | 1.45 | 4.55E-08 | 1.11E-06 | 1.13 | 1.29E-05 | 1.42E-03 |
| **TRAF3** | ENSG00000131323 | 27.8 | 0.58 | 7.40E-03 | 4.39E-02 | 0.84 | 1.07E-04 | 7.82E-03 |
| **TRPS1** | ENSG00000104447 | 17.6 | 1.09 | 3.84E-04 | 3.66E-03 | 0.94 | 1.34E-03 | 4.98E-02 |
| **TSPAN13** | ENSG00000106537 | 5.8 | 6.96 | 5.27E-06 | 8.34E-05 | 5.08 | 1.82E-04 | 1.16E-02 |
| **TSPAN17** | ENSG00000048140 | 25.6 | 0.84 | 2.46E-03 | 1.77E-02 | 0.95 | 5.64E-04 | 2.70E-02 |
| **TTYH3** | ENSG00000136295 | 5.3 | 2.36 | 6.21E-05 | 7.51E-04 | 1.80 | 9.18E-04 | 3.84E-02 |
| **U62317.1** | ENSG00000272666 | 5.2 | 3.30 | 5.54E-07 | 1.07E-05 | 1.82 | 9.99E-04 | 4.04E-02 |
| **UBASH3B** | ENSG00000154127 | 30.9 | 1.24 | 1.72E-06 | 3.03E-05 | 1.17 | 3.10E-06 | 4.28E-04 |
| **VDR** | ENSG00000111424 | 67.0 | 1.87 | 2.58E-13 | 1.73E-11 | 1.92 | 3.81E-14 | 8.05E-11 |
| **VMP1** | ENSG00000062716 | 123.1 | 0.53 | 9.93E-05 | 1.13E-03 | 0.56 | 3.13E-05 | 2.88E-03 |
| **ZBED2** | ENSG00000177494 | 5.9 | 5.64 | 8.61E-11 | 3.54E-09 | 3.31 | 2.96E-10 | 2.16E-07 |
| **ZBTB32** | ENSG00000011590 | 5.6 | 2.22 | 7.03E-04 | 6.13E-03 | 3.74 | 3.04E-07 | 5.74E-05 |
| **ZNF282** | ENSG00000170265 | 34.4 | 2.44 | 4.55E-10 | 1.63E-08 | 2.04 | 8.74E-08 | 2.25E-05 |
| **ZNRF1** | ENSG00000186187 | 29.0 | 2.72 | 6.38E-15 | 5.90E-13 | 1.37 | 1.31E-05 | 1.43E-03 |
| **uTreg core DOWN** | | | **DOWN vs bTreg** | | | **DOWN vs uTconv** | | |
| *Gene* | *Ensembl* | *baseMean* | *Log2(FC)* | *Pvalue* | *Padj* | *Log2(FC)* | *Pvalue* | *Padj* |
| **ABLIM1** | ENSG00000099204 | 131.6 | -2.07 | 5.82E-32 | 3.36E-29 | -0.84 | 3.61E-06 | 4.70E-04 |
| **AL157935.1** | ENSG00000227218 | 19.5 | -1.55 | 4.81E-08 | 1.16E-06 | -1.00 | 5.53E-04 | 2.66E-02 |
| **APBA2** | ENSG00000034053 | 17.8 | -1.10 | 1.54E-03 | 1.20E-02 | -1.44 | 1.55E-05 | 1.64E-03 |
| **ATF7IP2** | ENSG00000166669 | 56.4 | -1.44 | 1.84E-12 | 1.02E-10 | -0.95 | 3.88E-06 | 4.97E-04 |
| **BEX2** | ENSG00000133134 | 42.6 | -2.54 | 4.66E-20 | 8.90E-18 | -1.20 | 3.25E-05 | 2.96E-03 |
| **CCR7** | ENSG00000126353 | 605.6 | -1.83 | 2.50E-13 | 1.68E-11 | -0.94 | 1.80E-04 | 1.15E-02 |
| **GCSAM** | ENSG00000174500 | 29.7 | -1.25 | 4.56E-04 | 4.25E-03 | -1.96 | 8.19E-09 | 3.39E-06 |
| **GIMAP4** | ENSG00000133574 | 251.1 | -1.63 | 3.65E-23 | 1.07E-20 | -0.81 | 1.02E-06 | 1.65E-04 |
| **GIMAP7** | ENSG00000179144 | 352.4 | -2.13 | 1.31E-21 | 3.20E-19 | -0.74 | 1.01E-03 | 4.06E-02 |
| **IL7R** | ENSG00000168685 | 1201.0 | -0.71 | 8.53E-03 | 4.92E-02 | -1.62 | 1.27E-09 | 7.65E-07 |
| **ITGA6** | ENSG00000091409 | 53.7 | -2.28 | 4.53E-20 | 8.72E-18 | -0.90 | 5.64E-04 | 2.70E-02 |
| **LDLRAP1** | ENSG00000157978 | 166.2 | -2.14 | 8.14E-30 | 4.19E-27 | -0.62 | 1.32E-03 | 4.91E-02 |
| **LEF1** | ENSG00000138795 | 241.9 | -3.31 | 2.49E-57 | 4.22E-54 | -0.77 | 3.16E-04 | 1.78E-02 |
| **LINC02273** | ENSG00000245954 | 42.9 | -1.82 | 2.72E-08 | 6.96E-07 | -1.81 | 2.20E-08 | 7.49E-06 |
| **MGAT4A** | ENSG00000071073 | 100.1 | -1.09 | 5.05E-09 | 1.48E-07 | -1.07 | 5.30E-09 | 2.38E-06 |
| **PLAC8** | ENSG00000145287 | 136.7 | -1.02 | 1.12E-06 | 2.04E-05 | -1.09 | 1.17E-07 | 2.78E-05 |
| **PRKCB** | ENSG00000166501 | 114.2 | -1.23 | 1.27E-12 | 7.30E-11 | -0.85 | 1.19E-06 | 1.86E-04 |
| **RARRES3** | ENSG00000133321 | 208.5 | -0.62 | 4.25E-04 | 4.00E-03 | -0.70 | 5.82E-05 | 4.67E-03 |
| **RBL2** | ENSG00000103479 | 105.8 | -0.96 | 7.38E-08 | 1.71E-06 | -0.64 | 3.14E-04 | 1.78E-02 |
| **SATB1** | ENSG00000182568 | 163.1 | -0.80 | 1.10E-03 | 8.91E-03 | -1.26 | 2.07E-07 | 4.34E-05 |
| **TCF7** | ENSG00000081059 | 493.4 | -1.75 | 1.84E-25 | 6.80E-23 | -1.11 | 4.86E-11 | 4.28E-08 |
| **TTC39C** | ENSG00000168234 | 158.1 | -1.36 | 6.25E-25 | 2.18E-22 | -0.71 | 1.30E-07 | 2.98E-05 |
| **TTC9** | ENSG00000133985 | 40.7 | -0.99 | 8.71E-04 | 7.37E-03 | -1.15 | 9.23E-05 | 7.02E-03 |
