## Supplementary table 5 for "Human regulatory T cells at the maternal-fetal interface show functional site-specific adaptation with tumor-infiltrating-like features"

| **Supplementary table 5. Overlap of tumor-infiltrating Treg signatures with significantly upregulated genes in uTregs vs bTregs (% of overlapping genes/genes in signature)** | | | | | | |
| --- | --- | --- | --- | --- | --- | --- |
| **Overlap between uTregs and tumor-infiltrating Treg signatures** | | | | | | |
| Tirosh et al | Zheng et al | De Simone et al. | Pacella et al. | Toker et al. | Magnuson et al. | Plitas et al. |
| *75/172 (44%)* | *225/401 (56%)* | *92/309 (30%)* | *91/211 (43%)* | *42/124 (34%)* | *70/108 (65%)* | *105/423 (25%)* |
| ACP5 | ACP5 | ACAA2 | ACP5 | ARHGEF12 | ANXA4 | ACSL4 |
| AGTRAP | ACSL4 | ACP5 | ACTG2 | BCL2L1 | ARHGEF12 | ACTG2 |
| ANXA2 | ACTN4 | ACSL4 | AKAP13 | CD80 | ATF3 | ADPRH |
| B4GALT1 | AKIRIN2 | ACTG2 | AKIRIN2 | COL5A1 | BATF | AKAP5 |
| BATF | APOBEC3C | ADPRH | APOBEC3C | CRADD | BCL2L1 | ASB2 |
| BIRC3 | APOBEC3G | AKAP5 | ARID5B | CREB3L2 | CAPG | ATP6V1A |
| BST2 | ARHGEF12 | ARHGEF12 | ATP6V1B2 | CTSC | CCR5 | ATP6V1C2 |
| C17orf49 | ARID5B | ARNTL2 | ATP6V1D | DPYSL2 | CD74 | AURKA |
| C3AR1 | ARPP19 | AURKA | B3GAT1 | GADD45A | CD80 | BCL2 |
| CAP1 | ASB2 | BATF | BCL2L1 | GCNT1 | CDKN1A | C3AR1 |
| CCND2 | ASXL2 | BCL2L1 | CAMK1 | HIP1 | CPD | CCND2 |
| CD79B | ATP1B1 | CD7 | CCDC6 | ICOS | CREB3L2 | CCRL2 |
| CLIC1 | ATP6V1C2 | CDK6 | CCND2 | IL1R1 | CREM | CD40LG |
| CMC2 | BATF | CEACAM1 | CD2 | IL1R2 | CTNNA1 | CD7 |
| COX17 | BCL2L1 | CGA | CD4 | IL2RA | CTSH | CD79B |
| CRADD | BIRC3 | COL9A2 | CD63 | KAT2B | CXCR3 | CD80 |
| CTLA4 | BST2 | CRADD | CD7 | KBTBD8 | CYFIP1 | CDK6 |
| CTSC | BTG3 | CREB3L2 | CD74 | KSR1 | DUSP4 | CGA |
| CXCR3 | C21orf91 | CSF1 | CD79B | LAPTM4B | ENTPD1 | CKS2 |
| CXCR6 | C3AR1 | CTLA4 | CDK2AP1 | LAYN | FAM126A | COL9A2 |
| DPYSL2 | CCL20 | CTSC | CEACAM1 | LRRC32 | FAM129A | CORO1C |
| EIF4A1 | CCND2 | DPYSL2 | CLIC1 | LTA | FNDC3B | COX17 |
| ENO1 | CCR1 | EML2 | COL5A1 | MAST4 | FURIN | CPXM1 |
| EPS15 | CCR5 | ENTPD1 | COL9A2 | MYO5A | GCNT1 | CRADD |
| ERI1 | CD2 | ERI1 | COMMD7 | NAMPT | GRN | CREB3L2 |
| ICOS | CD63 | ETV7 | CRADD | NCOA3 | HIF1A | CSF1 |
| IFI6 | CD7 | EVA1B | CREB3L2 | NETO2 | HIVEP3 | CTSC |
| IL1R2 | CD74 | FBXO45 | CST7 | PDGFA | ICOS | DPYSL2 |
| IL2RA | CD79B | FKBP1A | CTNNA1 | PHACTR2 | IKZF4 | DYNC1I2 |
| IL2RB | CD80 | FNDC3B | CTSC | PHTF2 | IL1R2 | EGR3 |
| IL2RG | CD82 | FUCA2 | CXCR6 | PTP4A3 | IL1RL1 | ENTPD1 |
| ISG15 | CD83 | GADD45A | DPYSL2 | SAT1 | IRAK2 | ETV7 |
| LGALS1 | CDIP1 | GCNT1 | ENO1 | SEC14L1 | IRF5 | FAM126A |
| LGALS3 | CDKN1A | HAP1 | ENTPD1 | SGMS1 | ISG15 | FNDC3B |
| LY6E | CDKN2A | HAVCR2 | ERI1 | SLAMF1 | LAMP2 | FUCA2 |
| MAPKAPK3 | CEACAM1 | HIVEP3 | FBLN7 | SLC16A1 | LAPTM4B | GALM |
| MRPS6 | CLIC1 | ICOS | GALM | SOX4 | MAP2K3 | GCNT1 |
| MX1 | CMTM6 | IGFLR1 | GAPDH | TNFRSF13B | MAPKAPK2 | GEM |
| MYL6 | COL9A2 | IKZF4 | GLA | TNFRSF1B | MAPKAPK3 | HAVCR2 |
| NAMPT | CREB3L2 | IL1R2 | GRAMD4 | TPP1 | MRPS6 | HIVEP3 |
| NCF4 | CREM | IL1RL1 | HLA-DRB4 | WLS | MVP | HMOX1 |
| NDUFV2 | CSF1 | IL2RA | HSP90AA1 | ZBTB32 | MXD1 | HSPA1A |
| NFKBIZ | CST7 | IL2RB | IFI27 |  | NAMPT | ICOS |
| OAS1 | CTLA4 | KAT2B | IFI35 |  | NCF4 | IFT27 |
| PELI1 | CTNNA1 | KLHDC7B | IL18R1 |  | NDRG1 | IKZF4 |
| PGK1 | CTSC | LAPTM4B | IL1R1 |  | NINJ1 | IL1A |
| PHTF2 | CTSD | LAX1 | IL1R2 |  | OSBPL3 | IL1R1 |
| PLP2 | CXCR3 | LAYN | IRF4 |  | PICALM | IL1R2 |
| PRDM1 | CXCR6 | LEPROT | ITGB2 |  | PLP2 | IL1RL1 |
| PRDX3 | DDIT4 | LTA | KAT2B |  | PMAIP1 | IL1RN |
| PRNP | DDX24 | MAP1LC3A | LAPTM4B |  | RHBDD2 | IL2RA |
| PTTG1 | DPYSL2 | MAST4 | LAX1 |  | RHOC | IL4R |
| RPS27L | DUSP2 | MGST2 | LAYN |  | SAMSN1 | IRAK2 |
| S100A4 | DUSP4 | MICAL2 | LGALS1 |  | SDC4 | IRF5 |
| S100A6 | DYNLL1 | MINPP1 | LTA |  | SEC14L1 | KSR1 |
| SAMSN1 | EDARADD | MREG | MAPK6 |  | SKAP2 | LAPTM4B |
| SAT1 | ENO1 | NAB1 | MFHAS1 |  | SNX9 | LAYN |
| SDC4 | ENTPD1 | NCF4 | MGST2 |  | SSH1 | MAP2K3 |
| SDF4 | ERI1 | NDFIP2 | MYO5A |  | TMBIM1 | MGST2 |
| SLAMF1 | ETV7 | NETO2 | MZB1 |  | TNFRSF18 | MICAL2 |
| SNX5 | FAM129A | NUSAP1 | NAB1 |  | TNFRSF1B | MIR155HG |
| SPPL2A | FBLN7 | PAM | NFIL3 |  | TNFRSF4 | MRPS6 |
| SQSTM1 | FKBP1A | PARD6G | PAM |  | TNFRSF8 | MYO1E |
| TANK | FOS | PDGFA | PMAIP1 |  | TNFRSF9 | NAGA |
| TFRC | FOSL2 | PRDX3 | PMVK |  | TNIP2 | NCF4 |
| TMED9 | FYCO1 | PTP4A3 | PRDX5 |  | TRAF1 | NEDD9 |
| TMEM173 | GABARAPL1 | PTTG1 | PRNP |  | TRAF3 | NFKB2 |
| TNFRSF18 | GADD45A | RBKS | PSTPIP1 |  | UEVLD | NR4A1 |
| TNFRSF1B | GADD45G | RDH10 | RGS1 |  | VIM | NTRK1 |
| TNFRSF4 | GALM | RYBP | RHOB |  | ZNRF1 | PARD6G |
| TPM4 | GAPDH | SECTM1 | SAMD9 |  |  | PARPBP |
| TPP1 | GCNT1 | SLC16A1 | SCD |  |  | PDGFA |
| TXN | GOLGA8A | SNAP47 | SDC4 |  |  | PDIA6 |
| TYMP | GPI | SOX4 | SETBP1 |  |  | PHLDA1 |
| VDR | GRN | SPATS2L | SFT2D1 |  |  | PIK3AP1 |
|  | GSTO1 | SSH1 | SH2D1A |  |  | PSEN1 |
|  | HAVCR2 | SYT11 | SH2D2A |  |  | PTP4A3 |
|  | HERPUD1 | TFRC | SLC20A1 |  |  | PTTG1 |
|  | HLA-DQB1 | THADA | SOCS1 |  |  | RBBP8 |
|  | HLA-DRB1 | TMEM184C | SPPL2A |  |  | RBKS |
|  | HLA-J | TNFRSF18 | TFRC |  |  | RHOC |
|  | HNRNPLL | TNFRSF4 | TMCO1 |  |  | SDC4 |
|  | HSPA1A | TNFRSF8 | TMEM109 |  |  | SETBP1 |
|  | HSPB1 | TNFRSF9 | TNFRSF1B |  |  | SH2D2A |
|  | ICOS | TOX2 | TNS3 |  |  | SLAMF1 |
|  | IFI6 | TPP1 | TPI1 |  |  | SLC4A2 |
|  | IGFLR1 | TRAF3 | TTYH3 |  |  | SLCO4A1 |
|  | IKZF4 | TSPAN17 | ZBED2 |  |  | SNX5 |
|  | IL10RB | VDR | ZBTB32 |  |  | SPATS2L |
|  | IL18R1 | YIPF6 | ZFAND5 |  |  | THADA |
|  | IL1R1 | ZBED2 | ZFP36L1 |  |  | TNFRSF18 |
|  | IL1R2 | ZNF282 |  |  |  | TNFRSF1B |
|  | IL2RA |  |  |  |  | TNFRSF4 |
|  | IL2RB |  |  |  |  | TNFRSF8 |
|  | IL4R |  |  |  |  | TNFRSF9 |
|  | IQGAP1 |  |  |  |  | TNIP2 |
|  | IRF5 |  |  |  |  | TNIP3 |
|  | ISG15 |  |  |  |  | TNS3 |
|  | IVNS1ABP |  |  |  |  | TRAF1 |
|  | KAT2B |  |  |  |  | TYMP |
|  | KDM5B |  |  |  |  | UBASH3B |
|  | KIF20B |  |  |  |  | VDR |
|  | LAPTM4A |  |  |  |  | VRK2 |
|  | LAPTM4B |  |  |  |  | XXYLT1 |
|  | LAT2 |  |  |  |  | ZBED2 |
|  | LAYN |  |  |  |  |  |
|  | LDHA |  |  |  |  |  |
|  | LEPROT |  |  |  |  |  |
|  | LGALS1 |  |  |  |  |  |
|  | LGALS3 |  |  |  |  |  |
|  | LINC00963 |  |  |  |  |  |
|  | LRPAP1 |  |  |  |  |  |
|  | LTA |  |  |  |  |  |
|  | LYST |  |  |  |  |  |
|  | MAF |  |  |  |  |  |
|  | MAP2K3 |  |  |  |  |  |
|  | MAPKAPK3 |  |  |  |  |  |
|  | MAST4 |  |  |  |  |  |
|  | MCL1 |  |  |  |  |  |
|  | MICAL2 |  |  |  |  |  |
|  | MIR497HG |  |  |  |  |  |
|  | MKNK1 |  |  |  |  |  |
|  | MRPS6 |  |  |  |  |  |
|  | MYO5A |  |  |  |  |  |
|  | MZB1 |  |  |  |  |  |
|  | NAMPT |  |  |  |  |  |
|  | NCF4 |  |  |  |  |  |
|  | NCOA3 |  |  |  |  |  |
|  | NDFIP2 |  |  |  |  |  |
|  | NEDD9 |  |  |  |  |  |
|  | NRBP1 |  |  |  |  |  |
|  | OAS1 |  |  |  |  |  |
|  | P2RY10 |  |  |  |  |  |
|  | PAM |  |  |  |  |  |
|  | PDCD1 |  |  |  |  |  |
|  | PDIA6 |  |  |  |  |  |
|  | PELI1 |  |  |  |  |  |
|  | PFKFB3 |  |  |  |  |  |
|  | PGK1 |  |  |  |  |  |
|  | PHACTR2 |  |  |  |  |  |
|  | PHLDA1 |  |  |  |  |  |
|  | PHPT1 |  |  |  |  |  |
|  | PHTF2 |  |  |  |  |  |
|  | PIM3 |  |  |  |  |  |
|  | PKM |  |  |  |  |  |
|  | PLTP |  |  |  |  |  |
|  | PMAIP1 |  |  |  |  |  |
|  | PMF1 |  |  |  |  |  |
|  | PMF1-BGLAP |  |  |  |  |  |
|  | PPP1CB |  |  |  |  |  |
|  | PRDM1 |  |  |  |  |  |
|  | PRDX1 |  |  |  |  |  |
|  | PRDX3 |  |  |  |  |  |
|  | PRDX5 |  |  |  |  |  |
|  | PRF1 |  |  |  |  |  |
|  | PRKAR1A |  |  |  |  |  |
|  | PRNP |  |  |  |  |  |
|  | PTP4A1 |  |  |  |  |  |
|  | PTP4A3 |  |  |  |  |  |
|  | PTPN22 |  |  |  |  |  |
|  | PTPN7 |  |  |  |  |  |
|  | PTTG1 |  |  |  |  |  |
|  | RAB10 |  |  |  |  |  |
|  | RAB11FIP1 |  |  |  |  |  |
|  | RAB8B |  |  |  |  |  |
|  | RALGDS |  |  |  |  |  |
|  | RBPJ |  |  |  |  |  |
|  | RGS1 |  |  |  |  |  |
|  | RHBDD2 |  |  |  |  |  |
|  | RHOC |  |  |  |  |  |
|  | RNF19A |  |  |  |  |  |
|  | RPS27L |  |  |  |  |  |
|  | SAMSN1 |  |  |  |  |  |
|  | SAT1 |  |  |  |  |  |
|  | SCO2 |  |  |  |  |  |
|  | SDC4 |  |  |  |  |  |
|  | SDF4 |  |  |  |  |  |
|  | SERPINE2 |  |  |  |  |  |
|  | SFT2D1 |  |  |  |  |  |
|  | SH2D2A |  |  |  |  |  |
|  | SKAP2 |  |  |  |  |  |
|  | SLA |  |  |  |  |  |
|  | SLAMF1 |  |  |  |  |  |
|  | SLC16A1 |  |  |  |  |  |
|  | SLC3A2 |  |  |  |  |  |
|  | SLC5A3 |  |  |  |  |  |
|  | SNX9 |  |  |  |  |  |
|  | SPATS2L |  |  |  |  |  |
|  | SPPL2A |  |  |  |  |  |
|  | SQSTM1 |  |  |  |  |  |
|  | SRA1 |  |  |  |  |  |
|  | SRGN |  |  |  |  |  |
|  | STAT3 |  |  |  |  |  |
|  | SURF4 |  |  |  |  |  |
|  | SYT11 |  |  |  |  |  |
|  | TANK |  |  |  |  |  |
|  | TFRC |  |  |  |  |  |
|  | THADA |  |  |  |  |  |
|  | TMCO1 |  |  |  |  |  |
|  | TMEM173 |  |  |  |  |  |
|  | TMEM50A |  |  |  |  |  |
|  | TMX1 |  |  |  |  |  |
|  | TNFAIP3 |  |  |  |  |  |
|  | TNFRSF13B |  |  |  |  |  |
|  | TNFRSF18 |  |  |  |  |  |
|  | TNFRSF1B |  |  |  |  |  |
|  | TNFRSF4 |  |  |  |  |  |
|  | TNFRSF9 |  |  |  |  |  |
|  | TNIP3 |  |  |  |  |  |
|  | TOX2 |  |  |  |  |  |
|  | TPI1 |  |  |  |  |  |
|  | TPM4 |  |  |  |  |  |
|  | TPP1 |  |  |  |  |  |
|  | TRAF1 |  |  |  |  |  |
|  | TRAF3 |  |  |  |  |  |
|  | TRPS1 |  |  |  |  |  |
|  | TSPAN13 |  |  |  |  |  |
|  | TSPYL2 |  |  |  |  |  |
|  | TYMP |  |  |  |  |  |
|  | UBASH3B |  |  |  |  |  |
|  | VDR |  |  |  |  |  |
|  | VMP1 |  |  |  |  |  |
|  | ZBED2 |  |  |  |  |  |
|  | ZBTB32 |  |  |  |  |  |
|  | ZFP36L1 |  |  |  |  |  |
