## Supplementary table 6 for "Human regulatory T cells at the maternal-fetal interface show functional site-specific adaptation with tumor-infiltrating-like features"

| **Supplementary table 6. Genes most often shared between 7 tumor-infiltrating Treg signatures** | | | | |
| --- | --- | --- | --- | --- |
| *Name* | *# Signatures shared* | *DE ^pb^uTreg vs bTreg?* | *DE ^pb^uTreg vs ^inc^Treg?* | *In uTreg core?* |
| IL1R2 | 7 | Yes | No | Yes |
| TNFRSF1B | 6 | Yes | Yes | Yes |
| CTSC | 6 | Yes | Yes | Yes |
| LAPTM4B | 6 | Yes | No | Yes |
| DPYSL2 | 6 | Yes | No | Yes |
| CREB3L2 | 6 | Yes | No | No |
| ICOS | 6 | Yes | No | No |
| CCR8 | 6 | No | No | No |
| CSF2RB | 6 | No | No | No |
| GLRX | 6 | No | No | No |
| MAGEH1 | 6 | No | No | No |
| ENTPD1 | 5 | Yes | Yes | Yes |
| IL2RA | 5 | Yes | Yes | Yes |
| NCF4 | 5 | Yes | Yes | Yes |
| SDC4 | 5 | Yes | Yes | Yes |
| TNFRSF4 | 5 | Yes | Yes | Yes |
| CRADD | 5 | Yes | No | Yes |
| LAYN | 5 | Yes | No | Yes |
| TNFRSF18 | 5 | Yes | No | Yes |
| BCL2L1 | 5 | Yes | Yes | No |
| GCNT1 | 5 | Yes | No | No |
| TIGIT | 5 | No | Yes | No |
| EBI3 | 5 | No | No | No |
| F5 | 5 | No | No | No |
| FOXP3 | 5 | No | No | No |
| GBP2 | 5 | No | No | No |
| IKZF2 | 5 | No | No | No |
| IL12RB2 | 5 | No | No | No |
| TBC1D8 | 5 | No | No | No |
| ACP5 | 4 | Yes | Yes | Yes |
| BATF | 4 | Yes | Yes | Yes |
| ERI1 | 4 | Yes | Yes | Yes |
| NAMPT | 4 | Yes | Yes | Yes |
| PTTG1 | 4 | Yes | Yes | Yes |
| TFRC | 4 | Yes | Yes | Yes |
| TPP1 | 4 | Yes | Yes | Yes |
| VDR | 4 | Yes | Yes | Yes |
| CD80 | 4 | Yes | No | Yes |
| COL9A2 | 4 | Yes | No | Yes |
| IKZF4 | 4 | Yes | No | Yes |
| IL1R1 | 4 | Yes | No | Yes |
| KAT2B | 4 | Yes | No | Yes |
| LTA | 4 | Yes | No | Yes |
| PTP4A3 | 4 | Yes | No | Yes |
| TNFRSF9 | 4 | Yes | No | Yes |
| ZBED2 | 4 | Yes | No | Yes |
| MRPS6 | 4 | Yes | Yes | No |
| SLAMF1 | 4 | Yes | Yes | No |
| ARHGEF12 | 4 | Yes | No | No |
| CCND2 | 4 | Yes | No | No |
| CD7 | 4 | Yes | No | No |
| CD79B | 4 | Yes | No | No |
| HTATIP2 | 4 | No | Yes | No |
| CARD16 | 4 | No | Yes | No |
| SIRPG | 4 | No | Yes | No |
| DFNB31 | 4 | No | No | No |
| DUSP16 | 4 | No | No | No |
| FCRL3 | 4 | No | No | No |
| FLVCR2 | 4 | No | No | No |
| GBP5 | 4 | No | No | No |
| MYO5C | 4 | No | No | No |
| SYNGR2 | 4 | No | No | No |
| TMPRSS6 | 4 | No | No | No |
| UGP2 | 4 | No | No | No |
